# GluDs are ionotropic dopamine receptors tuned by G-proteins

**DOI:** 10.64898/2026.05.10.723887

**Authors:** Haobo Wang, Mae G. Weaver, Elisa Carrillo, Iris Zheng, Jeffrey Khau, Anish Kumar Mondal, Nicholas Koylass, Anthony Yanez, Weiqi Gu, Zhaozhu Qiu, Evan S. O’Brien, Vasanthi Jayaraman, Edward C. Twomey

## Abstract

Dopamine is a neurotransmitter essential for cognition, and its dysregulation is associated with neurological diseases^1,2^. Historically, dopamine has been understood to signal exclusively through metabotropic receptors^3^. Delta-type ionotropic glutamate receptors (GluDs), which have recently been established as ligand-gated ion channels^4,5^, are fundamental for synaptic maintenance, are implicated in neurological disorders, and co-localize with dopaminergic machinery. Here, we report that dopamine is a direct agonist of GluDs, eliciting ionotropic activity, as evidenced by bilayer recordings, cryo-electron microscopy (cryo-EM), mutagenesis, and cellular recordings. Dopamine binds to the GluD ligand-binding domain, inducing clamshell closure and channel activation through a distinct molecular interface. G-proteins act as molecular switches to tightly tune GluD activity: Gβγ inhibits ligand-gating, while Gα relieves Gβγ inhibition. These findings expand dopaminergic signaling, uncover how G-proteins regulate GluD channel activity, and provide a framework for studying the cellular activity of GluDs.

---

Dopamine was discovered over a century ago, and has since been appreciated as a neurotransmitter that is essential for learning, motor control, and motivation^1,6^. Accordingly, the dysregulation of dopaminergic signaling is linked to Parkinson’s disease, schizophrenia, depression, attention deficit hyperactivity disorder, bipolar disorder, and addiction^2^. Unlike many other neurotransmitters, dopamine has not been found to directly activate ligand-gated ion channels in the nervous system^3^. Rather, dopamine has long been thought to act via volume transmission by activating G-protein-coupled receptors (GPCRs), which then stimulate second-messengers to modulate neuronal excitability and plasticity^6–9^. Despite this view, dopamine is concentrated into vesicles similarly to other ionotropic neurotransmitters and is released at high concentrations, suggesting the presence of synaptic-like signaling mechanisms^3,10,11^.

Intriguingly, dopamine release machinery co-localizes with delta-type ionotropic glutamate receptors (GluDs; GluD1 & GluD2) in the molecular layer of the cerebellum (GluD2) and the dopaminergic midbrain (GluD1)^12–27^. GluDs are responsible for slow, depolarizing currents and are ligand-gated ion channels that are unique amongst ionotropic glutamate receptors (iGluRs): rather than binding glutamate, they bind distinct neurotransmitters like D-serine and GABA^4,5,22,23,28–32^. GluDs have fundamental roles in regulating both excitatory and inhibitory synapses and are implicated in diseases such as schizophrenia, cerebellar ataxia, intellectual disability, dementia, and glioblastoma^30,33,34^. Notably, at Purkinje cell dendrites, where GluD2 is highly localized, dopamine induces a slow, depolarizing current, hinting at a direct functionality^13^.

GluDs are heavily regulated in cells by GPCRs: GPCR activation enables currents to be recorded from GluDs^22–25,28,35^, though the exact molecular mechanisms are unclear. Generally, GPCR activation leads to splitting of the inactive G-protein heterotrimer Gαβγ into Gα and Gβγ, which are released from the GPCR to stimulate downstream signaling cascades^36–38^. We hypothesize that one possibility underlying GluD activation is direct binding by G-proteins. Indeed, there is evidence that other canonical iGluR subtypes (Kainate and α-amino-3-hydroxy-5-methyl-4-isoxazolepropionic acid [AMPA]) directly bind G-protein subunits^39–44^. G-protein tuning of GluD ionotropic function would not be without precedent; inward rectifying potassium channels, voltage-gated calcium channels, and transient receptor potential (TRP) channels are directly regulated by G-proteins^45–54^.

Our ideas are twofold: (1) that dopamine is a ligand for GluDs, and (2) that G-proteins regulate GluDs directly by modulating their ligand-gating capabilities. To test this, we purified human GluD1 and GluD2. Dopamine elicits ligand-gated currents from GluDs in brain lipid bilayers in a concentration-dependent manner. Through cryo-EM of GluD2 in the presence of dopamine, we confirm that dopamine directly binds to the GluD2 ligand-binding domain (LBD), through a binding mechanism distinct from D-serine. G-protein binding to GluD2 directly regulates the ligand-gating activity of both dopamine and D-serine, and we observe through patch-clamp recording that ligands activate GluD2 in cells following G-protein stimulation. These data provide evidence for a potential expansion of dopamine signaling, redefine our understanding of how GluD channels are regulated, clarify *in situ* observations of GluD function, and provide avenues for therapeutic intervention of GluD signaling networks.

## Results

### Dopamine gating of GluDs

Extracellular dopamine spans a wide concentration range in the brain, from tonic levels of tens to hundreds of nanomolar^55–57^, to phasic transients reaching low micromolar^58,59^, and estimated peaks of micromolar to low millimolar near a release site^60^. To test if dopamine elicits ligand-gated responses from the GluDs, we first purified human GluD2 (Extended Data Fig. 1a) and reconstituted the receptors into brain lipid bilayers to test for ligand-gated activity.

We began by recording responses to D-serine, a known GluD2 ligand, from concentrations of 0.01 µM to 1 mM and observed responses to D-serine beginning at 5 µM, with a marked increase in apparent activity at 100 µM (Fig. 1a). These responses are characteristic of a ligand-gated ion channel, with stochastic transitions between closed (C) and open (O) conductance states (Fig. 1b).

**Fig. 1.**
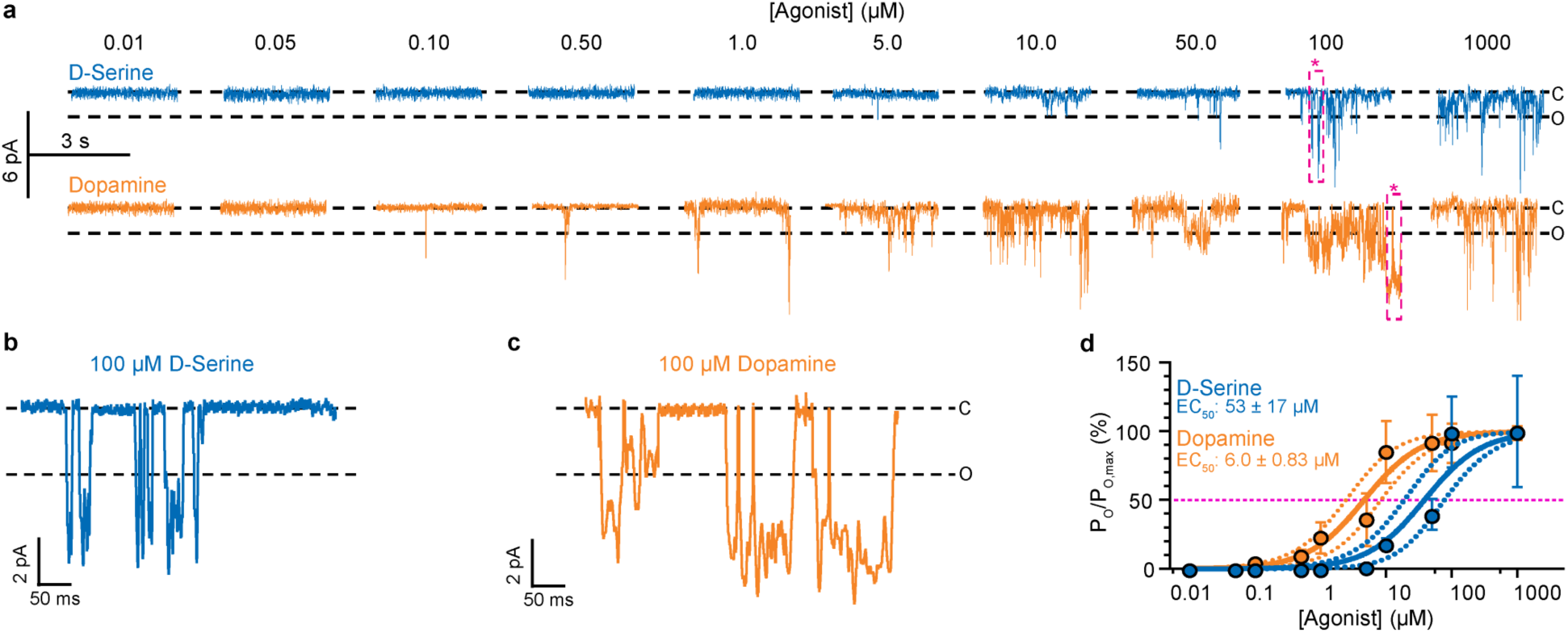
Ligand-gating of isolated GluD2 by dopamine. **a**, Example 3-s current traces of GluD2 in the presence of 0.01-1000 μM D-serine (*top*, blue) or dopamine (*bottom*, orange), highlighting the closed (C) and open (O) state conductance. **b,c,** 450 ms window of example current traces (purple box and asterisks, a) in the presence of 100 μM D-serine (b) or dopamine (c). **d,** The Hill equation (solid lines) ± the 95% CI (dotted lines) fitted to the D-serine and dopamine dose-response data (circles, mean ± s.e.m.) normalized to the P_O_ measured at 10,000 μM dopamine (Extended Data Fig. 1b, P_O_/P_O_,max). Dashed line, EC_50_. D-serine: EC_50_ = 53 ± 17 µM; 0.01 µM: 0.00 ± 0.00%, n = 5; 0.05 µM: 0.00 ± 0.00%, n = 5; 0.10 µM: 0.00 ± 0.00%, n = 5; 0.50 µM: 0.00 ± 0.00%, n = 5; 1 µM: 0.00 ± 0.00%, n = 5; 5 µM: 1.58 ± 1.58%, n = 6; 10 µM: 18.25 ± 3.89%, n = 7; 50 µM: 39.59 ± 11.28%, n = 7; 100 µM: 99.62 ± 25.87%, n = 7; 500 µM: 91.69 ± 40.33%, n = 5; 1000 µM: 100.0 ± 40.47%, n = 5. Dopamine: EC_50_ = 6.0 ± 0.83 µM; 0.01 µM: 0.00 ± 0.00%, n = 5; 0.05 µM: 0.30 ± 0.10%, n = 4; 0.10 µM: 3.97 ± 1.90%, n = 5; 0.50 µM: 8.96 ± 5.17%, n = 6; 1 µM: 22.57 ± 11.24%, n = 3; 5 µM: 35.66 ± 19.10%, n = 6; 10 µM: 84.99 ± 22.71%, n = 7; 50 µM: 91.53 ± 20.40%, n = 5; 100 µM: 91.39 ± 14.32%, n = 7; 500 µM: not determined; 1000 µM: 98.85 ± 5.04%, n = 13. All repeats are independent bilayer recordings.

Next, we tested whether dopamine elicited similar activity from GluD2 by performing dose-response recordings from 0.01 µM to 1 mM dopamine. Compared to D-serine, we observed currents beginning at 0.1 µM, with apparent activity increasing at 10 µM. Consistent with D-serine, GluD2 exhibits stochastic ligand-gating activity in the presence of dopamine (Fig. 1c). Thus, dopamine appears to be effective within tonic dopamine concentration ranges (e.g., 0.1 µM), where D-serine is less potent.

We calculated the relative potency for both D-serine and dopamine for GluD2 by measuring the open probability (P_O_) across the dose-responses. The half-effective concentration (EC_50_; ± standard error of the mean, s.e.m.) for D-serine was 53 ± 17 µM, while the EC_50_ for dopamine was 6.0 ± 0.83 µM, nine-fold more potent than D-serine while gating with equal efficacy (Extended Data Table 1). These data point to higher selectivity for dopamine versus D-serine in GluD2.

To verify that the dopaminergic currents were carried by GluD2, we tested whether the currents were blocked by 1-naphthyl acetyl spermine (NASPM), a polyamine-based toxin that blocks GluD currents^4,31,61,62^. Accordingly, we observed attenuation of currents from 10 mM dopamine in the presence of 100 µM NASPM, reflected by a decrease in both P_O_ and mean conductance (Extended Data Fig. 1b).

Next, we tested whether GluD1 was also gated by dopamine (Extended Data Fig. 1c). Like GluD2, we observed dopamine responses at 100 µM from GluD1. However, compared to GluD2, we observed lower P_O_ and mean conductance in GluD1 (Extended Data Fig. 1c). Importantly, 10 mM dopamine in the absence of purified GluDs, and GluDs in the absence of ligand did not show bilayer currents (Extended Data Fig. 1d). Therefore, we conclude that dopamine elicits ligand-gated channel activity across GluD subtypes in isolation.

To date, the canonical method of assaying GluD cellular function has been through recording spontaneous cellular currents from GluD2 Lurcher (Lc) mutants. Ligands such as D-serine reduce the constitutively active depolarizing currents, presumably through a desensitization-or inactivation-like mechanism. Therefore, we tested whether dopamine attenuates GluD2_Lc_ responses through cellular recordings from GluD2_Lc_. Indeed, we see a dopamine-dependent reduction of GluD2_Lc_ current, consistent with the reduction of depolarizing currents observed in D-serine (Extended Data Fig. 2). Of note is that the half-maximum inhibitory concentration (IC_50_) for dopamine inhibition of cellular GluD2_Lc_ is roughly 10^3^ times weaker (IC_50,Lc_ = 8.5 ± 2.0 mM) than the EC_50_ for dopamine activation of isolated, WT GluD2. Later experiments in this report detail the EC_50_ for dopamine activation of cellular WT GluD2.

### Dopamine binding to GluD2

To identify the dopamine binding site, we used cryo-EM to reconstruct the three-dimensional architecture of GluD2 in the presence of dopamine (Fig. 2a; Extended Data Figs. 3,4; Extended Data Table 2). The overall architecture of GluD2 was reminiscent of GluD2 bound to D-serine: the receptor has an overall italic “Y” shape in a non-domain-swapped arrangement, where local dimers in the amino-terminal domain (ATD) form between the A and B subunit positions, as well as the C and D subunit positions, and dimer pairings did not swap at the level of the LBD. Below the LBD is the transmembrane domain (TMD), which houses a central cation channel. The dopamine-bound GluD2 structure was similar to the D-serine-bound structure^4^ with an overall root mean square deviation (RMSD) of 1.2 Å, with each domain layer having an RMSD of 1.0 Å or less (Extended Data Fig. 5a).

**Fig. 2.**
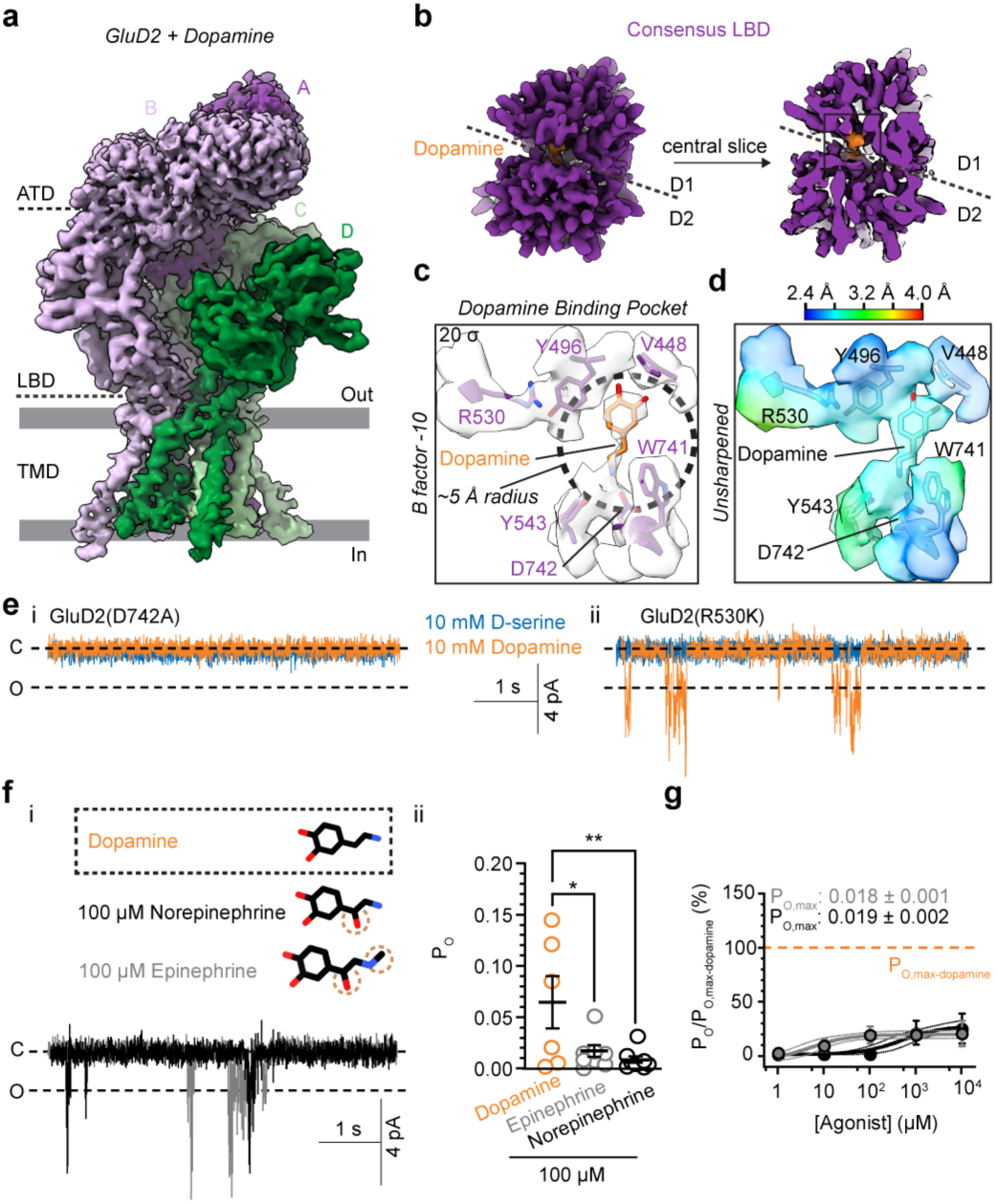
Cryo-EM of dopamine-bound GluD2. a, Unsharpened composite cryo-EM map of dopamine-bound GluD2. **b,** Structure and slice-through of a dopamine-bound LBD subunit monomer. **c,** Model of the LBD binding pocket (purple) bound to dopamine (orange) overlaid with the corresponding density at a contour threshold of 20 σ. **d,** Local resolution map computed for each voxel of the unsharpened dopamine binding pocket, FSC = 0.143. **e,** Example current traces of GluD2 (D742A; i) in the presence of 10 mM dopamine (n = 4, orange) or 10 mM D-serine (n = 4, blue) and of GluD2 (R530K; ii) in the presence of 10 mM dopamine (n = 4) or 10 mM D-serine (n = 10). **f,** Additional catecholamines tested (i), highlighting the differences from the dopamine structure (*top*) and example current traces in the presence of 100 µM norepinephrine (black) or epinephrine (grey) with associated P_O_ values (mean ± s.e.m.; ii). Dopamine: 0.065 ± 0.025, n = 6; norepinephrine: 0.009 ± 0.003, n = 9, P = 0.008, 95% CI of difference = 0.015 to 0.097, df = 20; epinephrine: 0.017 ± 0.006, n = 8, P = 0.03, 95% CI of difference = 0.005 to 0.0899, df = 20. ANOVA (F = 5.6, p = 0.011) with Dunnett’s correction relative to dopamine. **g,** The Hill equation (solid lines) ± the 95% CI (dotted lines) fitted to the norepinephrine and epinephrine dose-response data (circles, mean ± s.e.m.), normalized to the P_O_ at 10,000 μM dopamine (dashed line, P_O_/P_O_,max-dopamine). Norepinephrine: EC_50_ = 500 ± 220 µM, P_O_,max = 0.019 ± 0.002; epinephrine: EC_50_ = 4.8 ± 1.7 µM, P_O_,max = 0.018 ± 0.001. All repeats are independent bilayer recordings.

All LBDs have closed clamshells (closed ∼16° compared to GluD2 in the absence of ligand), which is a general indicator of ligand-binding for iGluRs (Extended Data Fig. 5b)^63^. To unequivocally define the binding site, we used symmetry expansion to average the LBDs together (Extended Data Figs. 3,4; Extended Data Table 2), which resulted in a consensus LBD map at 2.5 Å resolution (Fig. 2b). This enabled placement of dopamine in the LBD, based on an unmodeled planar feature reminiscent of a catechol ring with a tail in a hydrophobic pocket (Fig. 2c) with a relative local resolution of ∼2.4 to 3.0 Å at this site (Fig. 2d). Of note, around the dopamine binding site (within a 5 Å radius of dopamine) are D742, W741, Y543, Y496, and V448. D742 is a critical residue for coordination of D-serine in the GluD2 LBD^4,31^. An important distinction, however, from the D-serine binding site is that R530, which is critical for D-serine coordination, is exterior from the dopamine site (Fig. 2c).

Thus, D742 and R530 are two landmark residues for testing the dopamine binding site versus D-serine; D742 is a charged moiety that coordinates both dopamine and D-serine, but only D-serine is coordinated by R530^4,64^. First, we tested the ligand-gating of GluD2(D742A), which expectedly, ablated bilayer currents in the presence of 10 mM D-serine, or 10 mM dopamine (Fig. 2e, inset i). Next, we recorded from GluD2(R530K) in the presence of D-serine or dopamine. While GluD2(R530K) ablated D-serine ligand-gating, it did not appear to readily affect ligand-gating by dopamine (Fig. 2e), consistent with our model, where R530K is > 5Å from the dopamine binding site (Fig. 2c) and shielded from dopamine by Y496 (Extended Data Fig. 5d).

Next, we tested the specificity of the GluD2 LBD for dopamine versus two other neuronal catecholamines: norepinephrine and epinephrine. Compared to dopamine, norepinephrine contains an additional hydroxyl group on the ethylamine side chain; epinephrine contains both this hydroxyl group and a methylated amine group (Fig. 2f, inset i). We observed GluD2 currents elicited by both 100 µM norepinephrine and epinephrine (Fig. 2f, inset ii; Extended Data Table 1). To further assess these catecholamines, we extended our recordings of norepinephrine and epinephrine across a dose-response (1 to 10,000 µM) (Fig. 2g). Strikingly, for both norepinephrine and epinephrine, the fitted maximal P_O_ reached was ∼75% lower than that of dopamine, suggesting that they act as partial GluD2 agonists (Extended Data Table 1). Collectively, these data suggest that the dopamine binding site is distinct from that of D-serine and preferentially sensitive to dopamine, with other catecholamines producing less efficacious ligand-gating.

### In vitro reconstitution of GluD2 regulation by G-proteins

GPCR activation stimulates currents from GluDs^22–25,28^. However, the molecular mechanisms of how cellular GluD currents are stimulated by GPCR activation are unknown. We hypothesized that activated Gα subunits may regulate GluD activation, considering the GTP-dependence of GluD currents^28^ and the evidence for direct Gα binding to other iGluRs^43,65^. Toward this end, we first tested whether G-proteins can directly bind GluDs, focusing on the most predominant family of G-proteins in the brain, the G_i/o_ inhibitory family^66^.

We directly tested if GluD2 can form complexes with G-proteins by using purified recombinant GluD2 immobilized on affinity resin as bait to assay GluD2 binding to purified Gα, Gβγ, and Gαβγ (Fig. 3a, inset i; Extended Data Fig. 6). We observed that Gα, Gβγ, and Gαβγ were independently pulled down in a GluD2-specific manner (Fig. 3a, inset ii). We then tested whether Gα can bind GluD2 independently of Gβγ by assaying whether GluD2-Gβγ can pull down Gα (pre-incubated with 1 mM GTPγS to prevent heterotrimer formation). We observed binding (Fig. 3a, inset ii), indicating that Gα and Gβγ can simultaneously and independently bind GluD2 in the GluD2-Gβγ+Gα complexes. While both Gα and Gβγ were capable of binding at the same time, qualitatively, Gα appeared to bind better than Gβγ; G-protein heterotrimer in the presence of GDP resulted in nearly equal amounts of the two species, but when added separately, Gα appeared to dominate (Fig. 3a, inset ii).

**Fig. 3.**
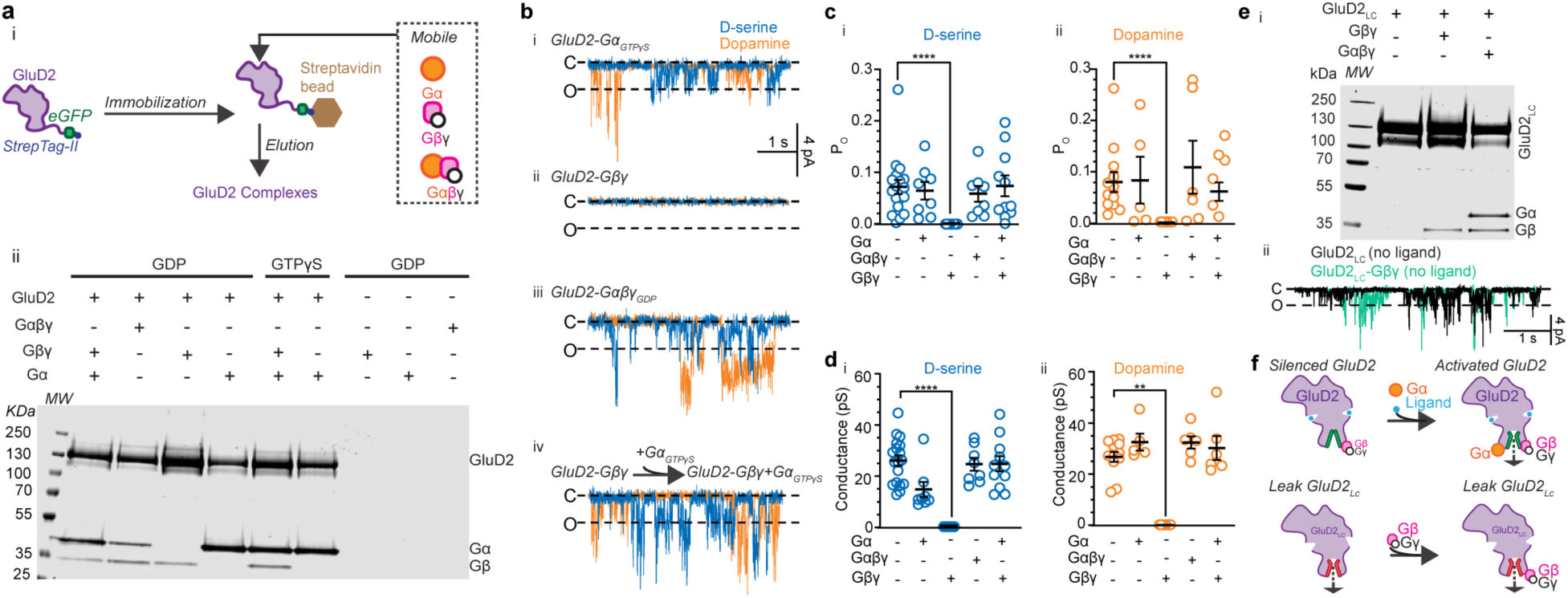
Reconstitution of GluD2 G-protein complexes. a, Schematic of G-protein affinity pull-down by GluD2-eGFP-StrepTag II (i) and example gel of the elutions (ii; n = 3). **b,** Example current traces in the presence of 10 mM D-serine (blue) or 10 mM dopamine (orange) of GluD2 bound to Gα (i; n = 8 and 5 traces for D-serine and dopamine, respectively), Gβγ (ii, n = 14 and 10), Gαβγ (iii, n = 8 and 6), and Gβγ rescue by Gα (iv, n = 11 and 6). **c**, Mean P_O_ ± s.e.m. of G-protein-free GluD2 currents in D-serine (i): 0.070 ± 0.013, n = 18 or dopamine (ii): 0.079 ± 0.020, n = 12. GluD2-Gα, D-serine: 0.063 ± 0.017, P > 0.99, mean rank difference = 1.19, Z = 0.16; dopamine: 0.082 ± 0.046, P > 0.99, mean rank difference = 3.57, Z = 0.59. GluD2-Gβγ, D-serine: 0.00 ± 0.00, P < 0.00010, mean rank difference = 30.44, Z = 5.0; dopamine: 0.00 ± 0.00, P < 0.00010, mean rank difference = 20.67, Z = 4.3. GluD2-Gαβγ, D-serine: 0.057 ± 0.015, P > 0.99, mean rank difference = 3.07, Z = 0.42; dopamine: 0.11 ± 0.052, P > 0.99, mean rank difference = 0.83, Z = 0.15. GluD2-Gα-Gβγ, D-serine: 0.072 ± 0.020, P > 0.99, mean rank difference = 0.76, Z = 0.12; dopamine: 0.061 ± 0.018, P > 0.99, mean rank difference = 1.8, Z = 0.32. Kruskal-Wallis test (Kruskal-Wallis statistic = 32 [D-serine], = 23 [dopamine]; P < 0.00010 [D-serine], = 0.00020 [dopamine]) with Dunn’s correction relative to G-protein-free GluD2. **d**, Mean conductance ± s.e.m. of G-protein-free GluD2 currents in the presence of D-serine (i): 25.44 ± 2.08 pS, n = 18 or dopamine (ii): 26.75 ± 1.99 pS, n = 12. GluD2-Gα, D-serine: 14.17 ± 3.00 pS, P = 0.07, mean rank difference = 17.21, Z = 2.3; dopamine: 32.51 ± 3.23 pS, P > 0.99, mean rank difference =-5.22, Z = 0.87. GluD2-Gβγ, D-serine: 0.00 ± 0.00 pS, P < 0.00010, mean rank difference = 33.33, Z = 5.5; dopamine: 0.00 ± 0.00 pS, P = 0.0020, mean rank difference = 17.08, Z = 3.5. GluD2-Gαβγ, D-serine: 24.02 ± 2.51 pS, P > 0.99, mean rank difference = 1.21, Z = 0.17; dopamine: 32.39 ± 2.34 pS, P = 0.88, mean rank difference =-6.92, Z = 1.2. GluD2-Gα-Gβγ, D-serine: 24.19 ± 2.99 pS, P > 0.99, mean rank difference = 2.29, Z = 0.35; dopamine: 30.22 ± 4.68 pS, P > 0.99, mean rank difference =-0.42, Z = 0.074. Kruskal-Wallis test (Kruskal-Wallis statistic = 38 [D-serine], = 24 [dopamine], P < 0.00010 [D-serine], < 0.00010 [dopamine]) with Dunn’s correction relative to G-protein-free. **e,** Example gel of G-protein affinity pull-down by GluD2_Lc_-eGFP-StrepTag II (i; n = 3) and example GluD2_Lc_ (n = 4) and GluD2_Lc_-Gβγ (n = 3) current traces (ii) in the absence of ligand. **f**, Proposed mechanism for G-protein modulation of GluDs. All repeats are independent bilayer recordings.

To assess the functional effects of G-proteins on GluD2 function, we reconstituted GluD2-G-protein complexes into brain lipid bilayers for bilayer recordings in either supersaturating (10 mM) D-serine or dopamine. In the absence of G-proteins, GluD2 had a P_O_ of 0.070 ± 0.013 and 0.079 ± 0.020 in the presence of D-serine and dopamine, respectively (Fig. 3c). For GluD2-Gα and GluD2-Gαβγ, these values were unchanged: 0.063 ± 0.017 and 0.082 ± 0.046 (GluD2-Gα); 0.057 ± 0.015 and 0.11 ± 0.052 (GluD2-Gaβγ) in the presence of D-serine and dopamine, respectively. Unexpectedly, Gβγ silenced the GluD2 ionotropic response regardless of ligand, evidenced by the lack of observable currents (Fig. 3d).

We next tested whether Gα can rescue the silencing of GluD2-Gβγ complexes. First, we pulled down Gα with GluD2 along with Gβγ (Fig. 3a, inset ii). We then recorded from GluD2-Gβγ complexes and tested whether silenced GluD2-Gβγ in bilayers could be rescued by the addition of Gα in the presence of GTPγS. Indeed, both D-serine and dopamine elicited ionotropic behavior from GluD2-Gβγ+Gα_GTPγS_, suggesting that Gα can relieve inhibition by Gβγ (Fig. 3b, inset iv). Importantly, recovery of GluD2-Gβγ by Gα restored both the P_O_ and mean conductance to G-protein-free levels, regardless of ligand (Fig. 3c,d).

To further investigate the interaction of G-proteins with GluDs, we used cryo-EM to reconstruct the ultrastructure of the GluD2-Gαβγ complex to approximately 9 Å (Extended Data Fig. 7a,b). We observed that when in complex with Gαβγ, GluD2 adopted a conformation that has only been previously observed for the gain-of-function cerebellar ataxia mutation GluD2(R710W)^4^, such that the ATD and LBD of GluD2 rearranged into a symmetric dimer-of-dimers (Extended Data Fig. 7c,d), marked by 0.88 correlation coefficient, CC (Extended Data Fig. 7e,f), between the GluD2-Gαβγ and GluD2(R710W) extracellular domains. This provides some insights into how G-proteins may relieve Gβγ inhibition, but these exact mechanisms will require further investigation. However, we did observe a cytosolic-facing density that was not attributable to previous GluD2 maps^4^, where the extra density appeared to protrude into the micelle near the cytosolic-facing side of the TMD (Extended Data Fig. 7g). This suggests that the G-proteins may be directly interacting with the GluD2 cytosolic-facing TMD, and not the GluD2 C-terminal domain (CTD).

To test this idea, we truncated the CTD from GluD2 (GluD2_ΔCTD_) and tested its binding to G-proteins. Like WT, GluD2_ΔCTD_ formed a stable complex with G-proteins: Gα, Gβγ, and Gαβγ were independently pulled down by GluD2_ΔCTD_ (Extended Data Fig. 7h). We then reconstituted these GluD2_ΔCTD_-G-protein complexes in brain lipid bilayers to test if the CTD is required for G-protein modulation of GluD2 (Extended Data Fig. 7i). In the absence of G-proteins, GluD2_ΔCTD_ maintained ionotropic function to D-serine and dopamine, with no observable difference from WT (Extended Data Fig. 7j). Critically, GluD2_ΔCTD_ followed the same modulatory scheme as WT: ionotropic function when bound to Gα or Gαβγ, and silencing when bound to Gβγ, which was rescued by the addition Gα. This suggests that the G-protein regulatory interface is independent of the CTD, likely at the cytosolic interface of the TMD.

Collectively, these *in vitro* reconstitution results support a dynamic, modulatory role of G-proteins on GluD2 function, dependent on the specific G-protein assembly.

### Mechanism of the lurcher mutation

For decades, Lc mutations in GluD1 and GluD2 have been the standard for interrogating cellular GluD function. GluD2_Lc_ (GluD2[A654T]) has a missense mutation at the top of the ion channel helix at an essential point for channel gating^4^. This mutation decouples the LBD from the TMD, enabling ion channel opening in the absence of ligand^61^. Based on our data from WT, we hypothesized that GluD2_Lc_ has spontaneous channel activity due to uncoupling from channel silencing by Gβγ.

One potential idea for such a mechanism is that the GluD2_Lc_ does not interact with Gβγ. To test this, we purified human GluD2_Lc_ and tested whether it interacts with Gβγ. Surprisingly, GluD2_Lc_ pulled down Gβγ and Gαβγ, like WT GluD2 (Fig. 3e, inset i). Next, we tested whether GluD2_Lc_ is silenced by Gβγ by reconstituting GluD2_Lc_-Gβγ into brain lipid bilayers for ionotropic recordings. Apo GluD2_Lc_ behaved as expected in bilayers, showing leak currents in the absence of ligand (Fig. 3e, inset ii). However, we also observed qualitatively similar currents with GluD2_Lc_-Gβγ, suggesting that while GluD2_Lc_ can interact with Gβγ, it is not silenced by it like WT GluD2. This suggests that the Gβγ silencing observed for WT GluD2-Gβγ was not due to Gβγ preventing GluD2 insertion into lipid bilayers, as Gβγ complexes were unsilenced by Gα addition and were readily detected for GluD2_Lc_-Gβγ. Further, Gβγ was unable to silence GluD2_Lc_ function, where gating was already disrupted by the mutation.

We therefore conclude that GluDs are ligand-gated ion channels that exhibit canonical ligand-gating in isolation (Fig. 3f). Neurotransmitters such as dopamine, D-serine, and GABA^4,5,30^ can bind to the LBDs, which in turn open the M3 ion channel helices. Gβγ prevents the receptor from ligand-gating via binding at the TMD cytosolic interface, and inhibition can be rescued by Gα binding. Further, GluD2_Lc_ remains active even when Gβγ is bound, perhaps due to the activation network being already disrupted between the LBD and TMD, indicating that Gβγ requires an intact activation pathway to silence ligand-gating.

### G-protein activation enables cellular GluD currents

The data from our *in vitro* reconstitution suggests that G-proteins directly tune GluD2, and that silenced GluD2 can be activated by G-protein mobilization. Based on this, we hypothesized that G-protein activation *in situ* would enable GluD2 activation by ligands. This would be consistent with the observations that GPCR activation is required for GluD currents in brain tissue and in heterologous systems.

To test this, we expressed GluD2 in heterologous cells and recorded macroscopic currents via whole-cell patch-clamp electrophysiology. To stimulate G-protein activation, we included 100 µM GTPγS in the patch pipette. Uninfected cells did not exhibit activation in the presence of cellular GTPγS to external D-serine or dopamine (Extended Data Fig. 8a,b). Similarly, cells infected with GluD2 but without intracellular GTPγS did not exhibit GluD2 currents (Extended Data Fig. 8a,b), consistent with previous reports.

Cells expressing GluD2 with intracellular GTPγS exhibited marked currents in 1 mM D-serine that were reversibly blocked by 100 µM NASPM (Fig. 4a), reminiscent of the NASPM block observed in AMPARs^67^. We repeated these experiments with 100 µM dopamine, which elicited NASPM-blocked responses like those of D-serine (Fig. 4b). Interestingly, these currents appear non-desensitizing, with slow (∼50 ms) rise times (Extended Data Fig. 8), consistent with the slow, depolarizing currents attributed to GluD from slice recordings^13,23,25,28^. As expected, we did not observe currents from GluD2 with intracellular GTPγS in the presence of glutamate, which does not activate GluDs (Extended Data Fig. 8a)^4^.

**Fig. 4.**
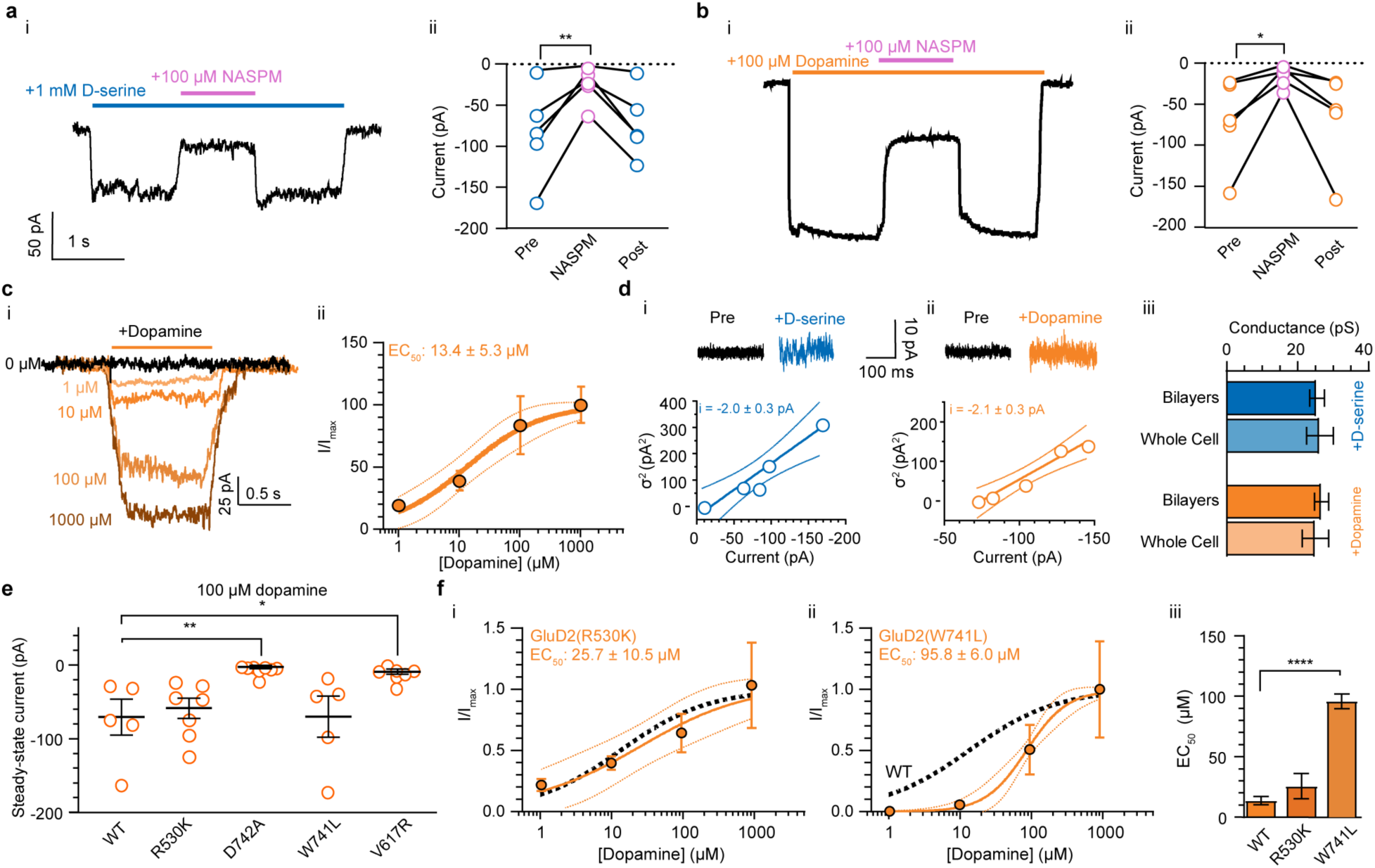
Cellular activation of GluD2. **a** Representative whole-cell traces (i) in response to D-serine (1 mM, blue bar) and the effects of NASPM (100 µM, pink bar) with the associated summary (ii) of steady-state current amplitudes elicited by D-serine before, during (P = 0.004, 95% CI of difference =-94.07 to-22.72, df = 8) and after (P = 0.68, 95% CI of difference =-45.88 to 25.47, df = 8) the application of NASPM (n = 5). Circles represent current amplitudes by cell. Repeated-measures ANOVA (F = 10.9, p = 0.005) with Dunnett’s correction relative to pre-NASPM. **b,** Representative whole-cell traces (i) in response to dopamine (100 µM, orange bar) and the effects of NASPM (100 µM, pink bar) with the associated summary (ii) of steady-state current amplitudes elicited by dopamine before, during (P = 0.02, 95% CI of difference =-98.40 to-8.107, df = 8) and after (P = 0.96, 95% CI of difference =-48.95 to 41.34, df = 8) the application of NASPM (n = 5). Repeated-measures ANOVA (F = 6.2, p = 0.02) with Dunnett’s correction relative to pre-NASPM. **c,** Representative whole-cell traces recording in response to various concentrations of dopamine (i) and the corresponding Hill equation (solid lines) ± the 95% CI (dotted lines) fitted to the steady-state current data normalized to 1000 μM (circles, mean ± s.e.m.). 0 μM (n = 20 cells), 1 μM (n = 6), 10 μM (n = 5), 100 μM (n = 5), 1000 μM (n = 5); df = 39, EC_50_ = 13.4 ± 5.3 µM. **d,** Representative traces (*top*) and summaries (*bottom*) of steady-state current variance (σ^2^) before and after the addition of 1 mM D-serine (i) or 100 μM dopamine (ii). Steady-state σ^2^-amplitude values (circles) by cell with a linear regression (solid line) ± the 95% CI (dotted lines). D-serine: 26.3 ± 3.8 pS, R^2^ = 0.93; dopamine: 25.0 ± 3.8 pS, R^2^ = 0.95. iii, σ^2^ analysis-derived conductance compared to bilayer-derived conductance (data from Fig. 1): D-serine: P = 0.99, 95% CI of difference =-10.35 to 11.96, df = 36; dopamine: P = 0.98, 95% CI of difference =-13.49 to 9.99, df = 36. ANOVA (F = 0.085, P = 0.97) with Tukey’s correction. **e**, Mean ± s.e.m. (lines) of steady-state macroscopic current amplitudes (circles) from GluD2 mutants (R530K, - 56.12 ± 14.48 pA, n = 7; D742A,-2.68 ± 2.40 pA, n = 8; W741L,-70.12 ± 28.00 pA, n = 5; V617R,-5.0 ± 1.57 pA, n = 6) compared to WT GluD2 (-70.52 ± 24.31 pA, n = 5). R530K, P > 0.99, mean rank difference =-1.31, Z = 0.25; D742A, P = 0.0030, mean rank difference =-17.48, Z = 3.4; W741L, P > 0.99, mean rank difference =-0.60, Z = 0.10; V617R, P = 0.046, mean rank difference =-13.93, Z = 2.5. Kruskal-Wallis test (Kruskal-Wallis statistic = 22, P = 0.00020) with Dunn’s correction relative to WT. **f**, Dopamine dose-response of steady-state current amplitudes of R530K (i) and W741L (ii). The Hill equation (solid lines) ± the 95% CI (dotted lines) fitted to the data (circles, mean ± s.e.m.) compared to the WT dose-response summary (black line). EC_50_ values ± s.e.m. (iii): R530K, 0 μM (n = 15), 1 μM (n = 4), 10 μM (n = 5), 100 μM (n = 7), 1000 μM (n = 5), EC_50_ = 25.75 ± 10.46 μM, P = 0.49, 95% CI of difference =-38.91 to 14.80, df = 17. W741L, 0 μM (n = 14), 1 μM (n = 4), 10 μM (n = 4), 100 μM (n = 5), 1000 μM (n = 5), EC_50_ = 95.79 ± 6.01 μM, P < 0.00010, 95% CI of difference =-98.65 to-65.55, df = 21. ANOVA (F = 37, P < 0.00010) with Dunnett’s correction relative to WT. All repeats are from individual, separate cells.

We measured a dose-response from 0 to 1000 µM dopamine with intracellular GTPγS (Fig. 4c, inset i) to estimate the relative potency of dopamine on cellular GluD2. We observed concentration-dependent steady-state current amplitudes that, when normalized to the maximum current response at 1000 µM, were well fit to the Hill equation (Fig. 4c, inset i; Extended Data Table 1). We estimated the EC_50_ of cellular GluD2 for dopamine to be 13.4 ± 5.3 µM (Fig. 4c, inset ii), not significantly different than the dopamine EC_50_ measured in bilayers (Fig. 1d; Extended Data Fig. 8c).

To cross-validate our cellular and bilayer recordings, we performed between-cell stationary variance (σ^2^) analysis of GluD2 currents elicited by 1 mM D-serine or 100 µM dopamine to assess unitary conductance. Consistent with previous reports and our measured P_O_, a linear regression model fit the σ^2^-current amplitude relationships for D-serine and dopamine well^23,25,28^. From the slope of these models, we calculated the unitary GluD2 conductance elicited by D-serine to be 26.3 ± 3.8 pS and dopamine to be 25.0 ± 3.8 pS (Fig. 4d, inset iii), consistent with our calculated bilayer values. Taken together, we conclude that G-protein activation in cells enables ligand-gating from GluD2 in a manner that is consistent with our bilayer experiments.

Next, we introduced point mutations in the LBD that we expect to be involved with dopamine entry and binding to the GluD2 LBD (D742A and W741L) or not involved (R530K) in dopamine coordination (Fig. 4e). As a negative control, we also tested GluD2(V617R), which has previously been shown to have reduced conductance^23,68^. Consistent with our bilayer data, we observed that GluD2(D742A), but not GluD2(R530K), significantly reduced the currents observed in 100 µM dopamine (Fig. 4e; Extended Data Fig. 8d), and GluD2(D742A) ablated currents completely. Further, we observed that GluD2(W741L), but not GluD2(R530K), reduced the potency of dopamine for GluD2, as measured by the EC_50_ (Fig. 4f; Extended Data Table 1).

To narrow down how cellular GluD2 ligand-gating is enabled by G-protein activation, we performed inside-out patch-clamp experiments, which enabled us to directly manipulate the cytosolic face of the channel and surrounding membrane. Expectedly, cellular GluD2 did not basally have ligand-gated responses to D-serine (Extended Data Fig. 9a, inset i) nor dopamine (Extended Data Fig. 9b, inset i). Next, on the same patch, we simulated GPCR activation with the Gβγ binding peptide (mSIRK) to stimulate Gα release from Gαβγ^69^. Correspondingly, we observed cationic currents after mSIRK treatment in the presence of D-serine or dopamine (Extended Data Fig. 9a,b, inset ii). We then applied NASPM, which attenuated the currents (Extended Data Fig. 9a,b, inset iii), as expected based on our bilayer and whole-cell recordings. We repeated these inside-out patch experiments in the presence of purified Gα. Akin to the mSIRK treatment, Gα enables GluD2 ionotropic behavior within cells in the presence of D-serine (Extended Data Fig. 9e, insets i-ii) and dopamine (Extended Data Fig. 9f, insets i-ii), and these currents were attenuated by NASPM (Extended Data Fig. 9e,f, inset iii). Finally, we confirmed that these currents were not elicited by glutamate, which does not activate GluDs (Extended Data Fig. 10).

These data, in combination with our *in vitro* reconstitution and whole-cell data, suggest that the GPCR-dependence of GluD currents^22–25,28^ is from activated G-proteins alleviating silencing by Gβγ (Figs. 3,4, Extended Data Figs. 8-10).

## Discussion

Dopamine has long been understood to not directly activate vertebrate ligand-gated ion channels^1–3^. However, here we show that dopamine directly elicits cationic currents via GluDs *in vitro* and in cells in a manner that is highly regulated by G-proteins. The dopamine-GluD connection may have previously been overlooked because cellular currents require G-protein activation, which likely liberates Gα to alleviate Gβγ silencing within the membrane microdomain; consistent with this, currents are not readily observed under basal conditions^70^.

G-proteins have multimodal effects on GluD signaling. A key future direction will be how the G-protein repertoire, including other alpha subunits such as the stimulatory (G_s_), G_q/11_, and G_12/13_ families, differentially tune GluD signaling across different stoichiometries, cellular contexts, neurotransmitter responses, and disease mutations.

Of interest is that GluD2 forms a complex with inactive Gαβγ. It has been suggested that GluDs may signal through metabotropic means by acting as *trans*-synaptic signal transduction devices^71^. One possibility is that *trans*-synaptic proteins such as the cerebellins and neurexins, which are tightly correlated with GluDs throughout the brain^33,34,72,73^, modulate the G-protein regulation of GluD channel activity.

A limitation of our work is that our *in vitro* reconstitution experiments are in saturating G-protein, isolated conditions that are likely distinct from the cellular milieu. What is clear, however, is that Gβγ silences GluD2, and Gα relieves the silencing. Higher resolution cryo-EM reconstructions will be required to elucidate the G-protein binding sites, and the G-protein:GluD2 stoichiometry relevant for physiological regulation.

The keys to unlocking the molecular function of GluDs are their G-protein regulators, which render GluDs as conditional ionotropic receptors. Understanding the depth of GluD regulation by G-proteins throughout brain tissue will be a key step to reclassifying the roles of GluDs in the future.

## Methods

### Construct design

The DNA coding sequence of full-length human GluD2 (UniProt ID O43424-1) was fused to a Thr-Gly-Gly linker, a thrombin cleavage site (LVPRGS), an enhanced GFP (eGFP) sequence, a Ser-Gly-Leu-Arg-Ser linker, a Strep-Tag II (WSHPQFEK) sequence, and a stop codon at its C-terminus. The construct was introduced into a pEG-BacMam vector for baculovirus-driven protein expression in HEK293 cells^74^. GluD2_Lc_ (A654T), GluD2(D742A), GluD2(W741L), GluD2(V617R), and GluD2(R530K) were generated by site-directed mutagenesis. GluD2_ΔCTD_ was generated by truncating amino acids 863-1007. The human Gα_i1_ sequence (Uniprot ID P63096) with an amino-terminal 6x His-tag, a HRV 3C protease cleavage site (LEVLFQGP), and a flexible glycine/serine linker were inserted into a pET21a plasmid (gifted by Dr. Daniel Hilger)^75^. Human Gβ₁γ₂ sequences (Uniprot ID P62873 for Gβ₁ and P59768 for Gγ₂) were used to express and purify the G-protein. The Gβ_1_ sequence was preceded by an N-terminal 6x His-tag, an HRV 3C protease cleavage site, and a Gly-Ser-Ser-Gly glycine/serine linker^75^.

### Expression and purification of human GluDs

Bacmids and baculoviruses of the full-length human GluD1, GluD2, and GluD2_Lc_ were made using standard methods^4^. In short, each cloned pEG-BacMam plasmid carrying a specific GluD2 construct was transformed into DH10Bac (Gibco, cat# 10301012) to generate bacmids. Those bacmids were then transfected into Expisf9 cells (Gibco, A35243) at a concentration of 2.5 million cells per mL using ExpiFectamine Sf transfection reagent (Gibco, A38915). The transfected cells were cultured at 27 °C with constant shaking. After 7 days, the supernatant of the cultured cells containing P1 baculoviruses was harvested and stored at 4 °C until further use. To increase the virus titer, P2 viruses were further gathered: 1 mL of the P1 viruses was added to 200 mL of fresh Expisf9 cells at a concentration of 1 million cells per mL. The infected cells were incubated at 27 °C with constant shaking. P2 viruses were isolated after 7 days via ultracentrifugation of the supernatant (41,300 xg for 60 minutes at 4 °C) and resuspended in 40 mL of Expi293media (ThermoFisher, A1435102). The concentrated P2 viruses were stored at 4 °C until further use.

For protein expression in mammalian cells, 20 mL of the concentrated P2 virus was added to Expi293F cells (Gibco, A14527) at ∼3 to 4 million cells per mL concentration. The infected Expi293F cells were incubated at 37 °C with 5% CO_2_ and constant shaking. At ∼16 hours post-infection, the cells were treated with 10 mM sodium butyrate (Sigma, 303410) and transferred to a 30 °C, 5% CO_2_ incubator. At 72 hours post-infection, the cells were harvested by centrifugation (500 xg for 20 minutes at 4 °C), washed with 1x PBS, and spun down again. The cell pellets were snap-frozen in liquid nitrogen and stored at-80 °C until further use.

For purification, the frozen cell pellets were resuspended at 4 °C in lysis buffer (20 mM Tris-HCl, pH 8.0, and 150 mM NaCl) with protease inhibitors (0.8 µM aprotinin, 2 µg/mL leupeptin, 2 µM pepstatin A, and 1 mM phenylmethylsulfonyl fluoride). The resuspended cells were then lysed on ice using a Misonix sonicator for four cycles (Fisher Scientific, 1s on and 1s off for 1 minute per cycle, power level 7). The supernatant was collected by centrifugation (2,500 xg for 20 minutes at 4 °C), and cell membranes were further isolated by ultracentrifugation (125,000 xg for 45 minutes at 4 °C). The pelleted membranes were then resuspended in lysis buffer (20 mM Tris-HCl, pH 8.0 and 150 mM NaCl) and mechanically homogenized before solubilized in solubilization buffer (20 mM Tris-HCl, pH 8.0 and 150 mM NaCl, 1% n-dodecyl-β-d-maltopyranoside (Anatrace, D310) and 0.2% brain extract total (Avanti Research, 131101P) as the final concentration) for 2 hours at 4 °C with constant stirring. Insoluble fractions were further removed by ultracentrifugation (125,000 xg for 45 minutes at 4 °C). The supernatant containing the solubilized membrane proteins was incubated with 2 mL of Strep-Tactin XT 4Flow resin (IBA, 2-5010) overnight at 4 °C with constant stirring. The following day, the resin was collected via a gravity column and washed with 10 column volumes (CVs) of glyco-diosgenin (GDN) buffer (150 mM NaCl, 20 mM Tris pH 8.0, and 0.01% GDN [Anatrace, GDN101]).

Bound proteins were eluted with GDN buffer containing 50 mM biotin (Thermo Scientific, PI29129) and concentrated down to 500 µL at 4 °C using a 100 kDa molecular weight cut-off concentrator (Millipore Sigma, UFC9100). The concentrated sample was then digested with thrombin (1:200 mass ratio of thrombin to eluted protein) for 1 hour at room temperature to remove the eGFP and StrepTag II. For the pull-down experiments, the thrombin digestion was skipped to preserve the tag. The protein samples (cleaved or uncleaved) were further separated via size-exclusion chromatography on a Superose 6 Increase 10/300 GL column (Cytiva, 29091596) equilibrated with the GDN buffer. The peak fractions corresponding to GluD2 tetramers were pooled and concentrated up to ∼4 mg/mL (Millipore Sigma, UFC5100) for cryo-EM specimen preparation. The samples were further ultracentrifuged to remove insoluble material after concentration. For the pull-down experiments, the protein sample was concentrated up to 8-10 mg/mL and snap-frozen in liquid nitrogen, stored in the-80 °C freezer until further use.

### Expression and purification of human G-proteins

The human Gα_i_ subunit was expressed and purified as previously described^75^. The pET21a plasmid carrying the Gα_i_ sequence was transformed into BL21(DE3) cells (NEB, C2527H). The cells were grown at 37 °C in Terrific Broth (Supelco, T0918) to an O.D.600 of 0.6, and protein expression was induced with 0.5 mM IPTG (Invitrogen, 15529019). The induced cells were incubated at 25 °C overnight with constant shaking (210 rpm) before being harvested and resuspended in lysis buffer (50 mM HEPES pH 7.5, 100 mM NaCl, 1 mM MgCl_2_, 50 μM GDP, 5 mM β-mercaptoethanol (βME), 5 mM imidazole, and protease inhibitors) the next day. The resuspended cells were lysed on ice by sonication using 50% power and 50% duty cycle for four times 60s. The supernatant was collected by centrifugation (2,500 xg for 20 minutes at 4 °C) and incubated with 5 mL of Talon resin (TaKaRa, 635503) for an hour. The bound resin was washed with 10 CV of the lysis buffer supplemented with 20 mM imidazole in a gravity column before the protein was eluted with the lysis buffer containing 250 mM imidazole. The eluted protein solution was dialyzed overnight in dialysis buffer (20 mM HEPES pH 7.5, 100 mM NaCl, 1 mM MgCl_2_, 20 μM GDP, 5 mM βME, and 5 mM imidazole). The 6x His-tag was cleaved by 3C protease (w/w 1:100), and 10 μL calf intestinal alkaline phosphatase (NEB, M0525S) was added to remove phosphorylation during the dialysis. The next day, uncleaved protein and 3C protease were removed by incubation with Talon resin for 1 hour at 4 °C. The flowthrough was concentrated down to 500 μL and run on a Superdex 200 Increase 10/300 GL column (Cytiva, 28990944) with SEC buffer (20 mM HEPES pH 7.5, 100 mM NaCl, 1 mM MgCl_2_, 20 μM GDP, and 100 μM TCEP). The corresponding fractions were pooled, concentrated, and stored at-80 °C until further use.

The human Gβ₁γ₂ heterodimer was expressed in Trichoplusia ni (High Five) insect cells (Gibco B85502) using a single baculovirus encoding the Gβ₁ and Gγ₂ sequences generated by the BestBac method (Expression Systems)^75,76^. The cells were infected at a density of 3-4 × 10^6^ cells per mL and incubated at 130 rpm and 27 °C for ∼60 hours. The cells were harvested by centrifugation (550 xg for 15 minutes at 4 °C), washed with 1x PBS, and spun down again. The cell pellets were snap-frozen in liquid nitrogen and stored at-80 °C until further use. The harvested cell pellets were resuspended in lysis buffer (10 mM Tris, pH 7.5, 5 mM βME, and protease inhibitors) for 1.5 hours. The membrane fraction was collected by ultracentrifugation (21,000 xg for 45 minutes at 4 °C) and homogenized manually using a glass Dounce homogenizer (Bellco Glass, 1984-10100) with solubilization buffer (20 mM HEPES pH 7.5, 100 mM NaCl, 1.0% sodium cholate, 0.05% DDM, 5 mM βME, and protease inhibitors). The solubilization reaction was further incubated at 4 °C for 1 hour with constant stirring. Insoluble debris was removed by ultracentrifugation (21,000 xg for 45 minutes at 4 °C). The supernatant was incubated with 4 mL talon resin for 1 hour at 4 °C before washing with 10 CV of the solubilization buffer supplemented with 20 mM imidazole. The bound protein was eluted with elution buffer (20 mM HEPES pH 7.5, 100 mM NaCl, 0.1% DDM, 5 mM βME, 250 mM imidazole). The eluted protein was treated with 3C protease (w/w 1:10) and 10 μL calf intestinal alkaline phosphatase (NEB, M0525S) and dialyzed overnight in 20 mM HEPES pH 7.5, 100 mM NaCl, 0.1% DDM, 5 mM βME, and 5 mM imidazole. The uncleaved protein and 3C protease were removed by incubating the overnight solution with 2 mL Talon resin. The flowthrough solution was collected and concentrated and run on a Superdex 200 Increase 10/300 GL column (Cytiva, 28990944) with SEC buffer (20 mM HEPES pH 7.5, 100 mM NaCl, 1 mM MgCl_2_, 20 μM GDP, and 100 μM TCEP). The corresponding fractions were pooled, concentrated and stored in-80 °C freezer until further use.

Human Gα_i1_β_1_γ_2_ heterotrimer was expressed and purified as previously described^75^. In brief, the heterotrimer was expressed and purified similarly to the Gβ₁γ₂ heterodimer, with the following differences. Two baculoviruses, one encoding the WT Gα_i1_ sequence and the other encoding the His-tagged Gβ₁ and Gγ₂ sequences, were generated by the BestBac method (Expression Systems). These viruses were used to co-transduce High Five insect cells (Gibco B85502) at a density of 3-4 × 10^6^ cells per mL and incubated at 130 rpm and 27 °C for ∼48-60 hours. During G-protein purification, all buffers were supplemented with GDP (10 μM) and MgCl_2_ (100 μM for lysis, 5 mM for solubilization, and 1 mM for dialysis and anion exchange buffers). Following reverse Ni affinity chromatography to remove uncleaved, His-tagged protein and 3C protease, the flowthrough solution, supplemented with 1 mM MnCl_2_, was collected and treated with CIP (NEB, M0525S), Antarctic phosphatase (NEB, M0289S), and λ phosphatase (NEB, P0753S) for 1 hour at 4 °C to dephosphorylate the protein. After adjusting the NaCl concentration to 50 mM by adding dilution buffer composed of 20 mM HEPES pH 7.4, 1 mM MgCl_2_, 10 μM GDP, 100 µM TCEP, and 0.05% DDM, the protein was loaded onto a Capto HiRes Q 10/100 (Cytiva) anion exchange column equilibrated in buffer A composed of 20 mM HEPES pH 7.4, 50 mM NaCl, 1 mM MgCl_2_, 10 μM GDP, 100 µM TCEP, and 0.05% DDM. The column was washed with buffer A, and bound protein was eluted over a linear gradient from 0 to 50% buffer B, which is near-identical to buffer A, but contains 1 M NaCl. The eluted lipidated G-protein heterodimer was concentrated using a 30-kDa MWCO Amicon spin concentrator (MilliporeSigma), flash frozen, and stored at-80°C until future use.

### Affinity pull-down of G-proteins

7 μM GluD2 (GluD2 or GluD2_Lc_) was incubated with 70 μM G-protein components (Gα_i_, Gβ₁γ₂, or Gα_i1_β_1_γ_2,_ 1:10 molar ratio) in a 50 μL reaction with buffer (20 mM HEPES pH 7.5, 100 mM NaCl, 1 mM MgCl_2_, 20 μM GDP, and 100 μM TCEP, 0.01% GDN) at 37 °C for 1 hour. Each reaction was then diluted to 200 μL in a 1 mL Eppendorf tube and incubated with 100 μL of Strep-Tactin resin at room temperature for 1 hour with constant mixing. The bound resin was washed three times with 200 μL buffer. For elution, the resin was incubated with 200 µL of reaction buffer supplemented with 50 mM biotin for 30 minutes, twice at room temperature. A total of 400 µL elution for each reaction was then pooled and concentrated using a 30K concentrator (Thermo Scientific, 88502) before running on a 4-20% Tris-glycine denaturing gel (ThermoFisher, WXP42012BOX). 1 mM GTPγS was added to the specific samples as indicated throughout the entire pull-down process (i.e., incubation, wash, and elution steps).

### Cryo-EM sample preparation and data collection

Homemade grids were used to prepare the cryo-EM samples in this study. In brief, C-flat holey carbon grids (Electron Microscopy Sciences, CF213-50-Au, Electron Microscopy Sciences) were first coated with 50 nm gold by a sputter coater (Leica EM ACE600) and then plasma cleaned using a Tergeo Plasma Cleaner (Pie Scientific) with Ar/O_2_ to make the final 1.2/1.3 gold grids with gold mesh, based on previously published methods^77,78^. The grids were glow-discharged for 120s with 25 mA current and 10s hold time in a Pelco Easiglow (Ted Pella, 91000) before sample application.

20 μL of the concentrated GluD2 sample was first pre-incubated in a thermocycler set at 37 °C for 10 minutes before use. Then, a 3 μL protein sample was spiked with 0.5 μL of the 70 mM dopamine (Sigma-Aldrich, H8502; final dopamine concentration, 10 mM), and mixed vigorously before being applied to freshly discharged grids in a FEI Vitrobot Mark IV (Thermo Fisher Scientific) chamber set at 37 °C and 90% humidity (wait time 30 s; blot force 5 N; blot time 4 s). The grids were then immediately plunge-frozen in liquid ethane.

For GluD2 dopamine, a total of 7,244 micrographs were collected on a 200-kV Glacios microscope (ThermoFisher Scientific) equipped with a Falcon 4i camera. The micrographs were collected with a total dose of 40.00 e^−^ / Å^2^, a dose rate of 9.93 e^−^ /pixel/s with a pixel size of 1.2 Å/pixel, and the defocus range was set from-0.5 μm to-2.5 μm.

For GluD2-Gαβγ, 50 µL of 7 µM purified GluD2 was mixed with 10 µL of 209 µM Gα_i1_β_1_γ_2_ heterotrimer (∼1:5 molar ratio) in a 200 µL total volume of the size-exclusion buffer (20 mM Tris, pH 8.0; 150 mM NaCl; 0.01% GDN) supplemented with 10 µM GDP and 1 mM MgCl_2_. The mixed solution was split into 4 PCR tubes and incubated in a thermocycler at 37 °C for 1.5 hours. After incubation, the samples were stored on ice before frozen on grids.

A 3 μL of protein sample was applied to freshly discharged homemade grids in a FEI Vitrobot Mark IV (Thermo Fisher Scientific) chamber set at 4 °C and 95% humidity (wait time 30 s; blot force 6 N; blot time 4 s). The grids were then immediately plunge-frozen in liquid ethane.

For GluD2-Gαβγ, a total of 2,657 micrographs were collected on a 200-kV Glacios microscope (ThermoFisher Scientific) equipped with a Falcon 4i camera. The micrographs were collected with a total dose of 40.00 e^−^ / Å^2^, a dose rate of 9.39 e^−^/pixel/s with a pixel size of 1.2 Å/pixel. The defocus range was set from-0.5 μm to-2.5 μm.

### Image processing

CryoSPARC^79^ (version 4.6.2) was used for processing sample images while final particle picking was performed with TOPAZ^80^. Details can be found in Extended Data Figs. 3,4.

### Model building, refinement, and structural analyses

ChimeraX^81^, Coot^82^, ISOLDE^83^ and PHENIX^84^ compiled by the SBgrid Consortium^85^ were used in combination to perform model building, refinement, and structural analysis. All visualizations and measurements were performed in ChimeraX. Model quality was assessed with MolProbity^86^.

### Bilayer recording and analyses

Unitary currents were recorded from purified GluD receptors using an Orbit mini with a temperature control unit (Nanion Technologies), following previously established methods^4,5^. Before all recordings, the Orbit mini was calibrated with a standard test cell chip (Nanion Technologies) to normalize baseline currents to 0 pA. A MECA recording chip (4x 100 µm cavities, Nanion Technologies, 132002) was then plugged into the Orbit mini and filled with recording solution (150 mM KCl, 20 mM HEPES, pH 7.4). A lipid bilayer with a capacitance of 10-25 pF was then formed over each recording cavity using a 10 µL pipette tip dipped in 100 mg/mL brain extract total (Avanti, 131101C) dissolved in decane (Sigma-Aldrich, D901-100ML). 1 µL of purified apo protein diluted to ∼4 µg/mL was added on top of each painted recording cavity. D-serine (ThermoFisher, A11353.14), dopamine hydrochloride (Sigma-Aldrich, H8502), (−)-Epinephrine (+)-bitartrate salt (Sigma-Aldrich, E4375) or L-(−)-Norepinephrine (+)-bitartrate salt monohydrate (Sigma-Aldrich, A9512) were then diluted in the recording solution to the noted concentrations. The working stocks of dopamine, epinephrine, and norepinephrine were made fresh from powder prior to every recording. During the recording, those stocks were stored on ice, covered with aluminum foil to minimize light exposure and oxidation. Every 30 minutes, the recording solution was discarded, and the recording chips were washed with blank buffer several times before a new batch of buffer containing one of the catecholamines was applied. The condition of the working stocks were visually inspected for color change, indicative of oxidation. In addition, catecholamine recordings were done in the dark except during the sample application and bilayer formation steps. To measure pharmacological block of dopaminergic currents, 100 µM NASPM (Tocris, 2766) was co-applied with dopamine. All recordings were conducted with a holding potential (V_hold_) of-80 mV and a maximum applied current of 200 pA at 37 °C. Currents were sampled at 20 kHz and written into digital files using the Elements Data Reader 4 (EDR4, Elements) software v.1.7.1. For each receptor subtype and each ligand, a separate MECA recording chip was used to prevent cross-contamination.

Currents were analyzed using Clampfit software version 11.3 (Molecular Devices). Currents were filtered with an 8-pole Bessel filter with a −3 dB cut-off frequency of 100 Hz and adjusted for baseline before analysis. These data were then idealized using the half-amplitude algorithm built into the Single-Channel Search function in Clampfit. Detection levels (<-2 pA) were predefined, and an automatic adjustment for baseline and simultaneous update for all levels function (with a 10% level contribution) function was used to correct for baseline drift. We omitted any event lasting less than 2 ms. The corresponding current amplitude values (i) and dwell times for each event were then used to calculate unitary point conductance and P_O_. P_O_ values were calculated using the total open dwell time divided by the overall length of the recording. Conductance (γ) was calculated by the following equation:

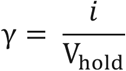

### Whole-cell recordings

For whole-cell experiments, currents were recorded from Expi293F cells at 48-72 hours post-infection with GluD2-eGFP-StrepTag II baculovirus. At ∼16 hours post-infection, the cells were treated with 10 mM sodium butyrate (Sigma, 303410) and transferred to a 30 °C, 5% CO_2_ incubator. On the day of recording, cells were added to 12-mm coverslips coated with 0.1 mg/mL poly-L-lysine and incubated at 37 °C with 5% CO_2_ for 5-20 minutes before recording. Whole-cell patch-clamp recordings were performed using pulled borosilicate glass pipettes (Sutter Instrument) of 2-6 MΩ resistance filled with intracellular solution containing (in mM) 135 CsF, 33 CsOH, 2 MgCl_2_, 10 HEPES, 11 EGTA, at pH 7.4 (CsOH), and were supplemented with 100 μM GTPγS as indicated. Extracellular solutions contained (in mM) 150 NaCl, 5 KCl, 2 CaCl_2_, 1 MgCl_2_, 10 HEPES at pH 7.4 (NaOH) and were supplemented with the indicated concentrations of D-serine, dopamine, or glutamate with or without 100 μM NASPM, as indicated. For dopamine experiments, oxidation was monitored by visual inspection of solutions, and solutions were changed every 30-60 minutes to prevent buildup of oxidized species.

Upon formation of the whole-cell configuration, the voltage was clamped at −80 mV, and cells were lifted and exposed to agonist-free extracellular solutions. Solution exchange of agonist-containing solutions was controlled using an SF-77C perfusion fast-step system (Warner Instruments), as previously described^87^. Agonist-containing solutions were applied for 0.5-1.5s and were followed by a 2s agonist-free wash. Recordings were amplified and low-pass filtered at 2 kHz using an Axopatch 700B amplifier (Molecular Devices), sampled at 2 kHz, and written into digital files with a Digidata 1550B digitizer (Molecular Devices) and pCLAMP 11 acquisition software.

To determine the apparent rates of activation and deactivation, we measured the 10-90% rise time of activation and fit a first-order Chebyshev exponential with a 4-pt smoothing filter to the offset of agonist-induced currents, respectively.

For the GluD2_Lc_ cellular recordings, mouse GluD2 (A654T) was transfected into HEK293T Cells with iMFectin at a 1:0.2 ratio per 35-mm dish. Patch pipettes (3–5 MΩ) were filled with an intracellular solution containing 135 mM CsF, 33 mM CsOH, 2 mM MgCl₂, 1 mM CaCl₂, 11 mM EGTA, and 10 mM HEPES. The extracellular solution contained 150 mM NaCl, 4 mM KCl, 2 mM CaCl₂, 10 mM HEPES, and dopamine (pH 7.4). External solutions were locally applied to lifted cells using a SF-77B perfusion fast-step (Warner Instruments). Whole-cell patch-clamp recordings were performed 24 h post-transfection at a holding potential of −60 mV. Currents were sampled at 10 kHz and low-pass filtered at 5 kHz using a 16-bit A/D converter (Digidata 1550A, Molecular Devices) controlled by pCLAMP 10 software (Molecular Devices).

### Stationary variance analysis

Stationary variance analysis was performed as previously described^25,89^. In brief, for each recording, we measured the variance (σ^2^) of steady-state current amplitudes (I), subtracting the baseline variance before agonist application. For each condition, we then plotted the normalized σ^2^ to cell-matched I. A linear regression was then fit to the σ^2^-I relationship, the slope of which is equal to the mean unitary current amplitude, which was then converted to unitary conductance.

### Inside-out recordings

HEK293T cells were maintained in Dulbecco’s modified Eagle’s medium (GenDEPOT) supplemented with 10% fetal bovine serum (GenDEPOT) and penicillin–streptomycin (Invitrogen). Cells were transiently transfected with GluD2 (human GluD2-eGFP-StrepTag II) using iMFectin Poly DNA Transfection Reagent (GenDEPOT) according to the manufacturer’s instructions.

Recordings were performed in the inside-out patch-clamp configuration. Patch pipettes (8–12 MΩ) were filled with an external solution containing 150 mM NaCl, 4 mM KCl, 2 mM CaCl₂, 10 mM HEPES, and either 10 mM D-serine or 10 mM dopamine (pH 7.4). The intracellular (bath) solution contained 135 mM CsF, 33 mM CsOH, 2 mM MgCl₂, 1 mM CaCl₂, 11 mM EGTA, and 10 mM HEPES. Recordings were performed at a holding potential of −100 mV. 25 µM mSIRK (Sigma-Aldrich, 371818), 70 µM Gi, and 300 µM NASPM were locally perfused around the patch. Currents were acquired at 50 kHz and low-pass filtered at 5–10 kHz using an Axopatch 200B amplifier and Digidata 1550A digitizer (Molecular Devices). For analysis, traces were further filtered at 1 kHz^88^.

Single-channel events were detected and analyzed in Clampfit (Molecular Devices) using the single-channel search function. Open and closed states were identified by amplitude threshold crossing, and traces were idealized accordingly. P_O_ was calculated as the total open time divided by the total recording duration.

## Statistical Analyses

All the data analyses were done using GraphPad Prism 11. Mean conductance and open probability were computed per bilayer, where n = the number of individual recorded traces per independently formed bilayer. Unless otherwise noted, all summary data represent mean ± s.e.m. Significance of difference was set to α = 0.05 and the normality of the data distribution was determined by the Shapiro-Wilk Test. For normally distributed data, a one-way ANOVA or Student’s t-test was used as noted. For non-normally distributed data, a Kruskal-Wallis test or Welch’s t-test was used. P values were corrected for multiple comparisons using either Tukey’s, Dunnett’s, or Dunn’s multiple comparisons test, as noted.

## Data Availability

The cryo-EM reconstructions are deposited into the Electron Microscopy Data Bank (EMDB). The whole maps are the primary cryo-EM maps in each deposition, and each local map, as applicable, and half maps are supplied as supplemental files in each deposition. All structural coordinates are deposited in the Protein Data Bank (PDB). The accession codes for each state (EMDB, PDB) will become available after publication.

## Acknowledgements

We thank the members of the Twomey and Jayaraman labs for insightful discussions during the development of this work. This work was supported by the National Institutes of Health (NIH; grants #R35GM154904, #R01NS148147), the One Mind Bristol Myers Squibb Rising Star Award, and the Lee Hood Prize in Biomedical Science via E.C.T, NIH grant #R35GM122528 to V.J., NIH grant #DP1DA066056 (E.S.O.), and NIH grants #RF1NS134549, # R35GM124824 to Z.Q. A.Y. was supported by NIH training grant #T32GM144272 and J.K. was supported by NIH training grant #T32GM135131. The cryo-EM data were collected at the Beckman Center for Cryo-EM at Johns Hopkins, which is supported by the Arnold and Mabel Beckman Foundation, the Howard Hughes Medical Institute, the Johns Hopkins University School of Medicine, and private, anonymous donors. All computational work was supported through the Johns Hopkins Research Information Technology DISCOVERY high-performance computing cluster.

## Author Contributions

E.C.T. conceptualized and supervised the project. H.W., M.G.W., V.J., and E.C.T. designed the experiments. H.W., M.G.W., and A.K.M. expressed and purified the GluD receptors. H.W. collected and analyzed the cryo-EM data. H.W. and E.C.T. built the molecular model. H.W., M.G.W., I.Z., and A.Y. collected the bilayer data. H.W. and M.G.W. performed the bilayer data analyses. H.W., J.K., and E.S.O. purified the G-proteins, with E.S.O. advising on G-protein complex experiment design. H.W., M.G.W., and I.Z. performed the G-protein complex experiments. E.C., M.G.W., W.G., V.J and E.C.T. designed the cellular electrophysiology experiments. M.G.W. and N.K. performed the whole-cell experiments, supervised by Z.Q. and E.C.T. M.G.W. analyzed the whole-cell electrophysiology data. E.C. performed the inside-out cellular electrophysiology experiments. E.C. and V.J. analyzed the inside-out electrophysiology data. H.W., M.G.W., and E.C.T. wrote the manuscript, which was edited by all authors.

## Additional Information

**Supplemental Information** The online version contains supplemental information.

**Correspondence and requests for material** should be addressed to Edward C. Twomey & Vasanthi Jayaraman.

## Extended Data Figures

**Extended Data Fig. 1.**
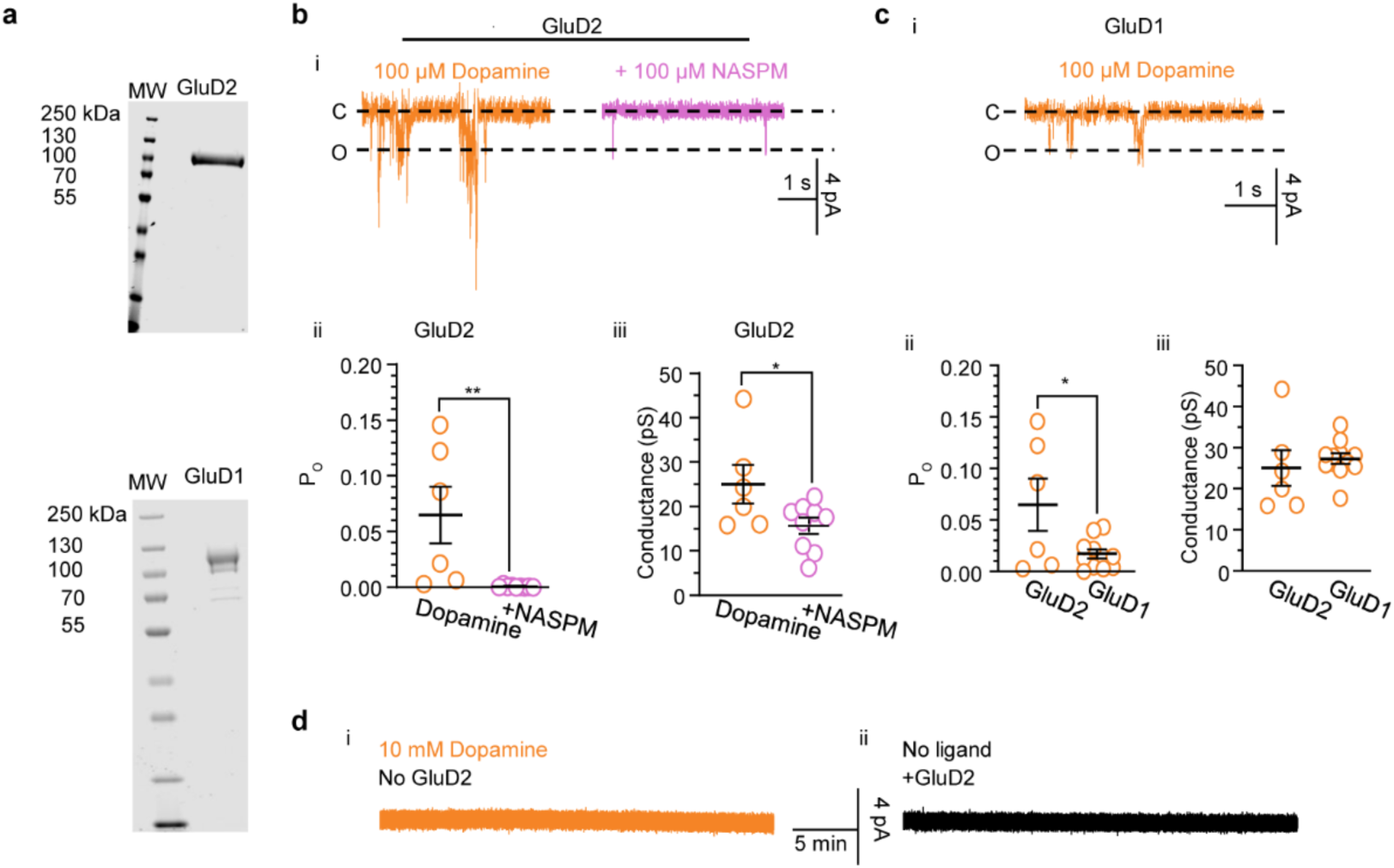
Comparison of dopaminergic activation of isolated GluD1 and GluD2. a, Example gels of purified GluD2 *(top)* and GluD1 *(bottom).* **b,** Example current traces (i) of GluD2 in response to 100 µM dopamine without (n = 6) or with (n = 9) the addition of 100 µM NASPM, with associated summaries (mean ± s.e.m.) of P_O_ (ii) and conductance (iii). ii, 100 µM dopamine: 0.065 ± 0.025; 100 µM dopamine + 100 µM NASPM: 0.00070 + 0.00030, t = 3.1, P = 0.0079, df = 13. iii, 100 µM dopamine: 25.02 + 4.37 pS, t = 2.2, P = 0.048, df = 13. Unpaired Student’s t-test. **c,** Example current traces (i) of GluD1 in response to 100 µM dopamine (n = 11) with associated summaries (mean ± s.e.m.) of P_O_ (ii) and conductance (iii). ii, 0.017 ± 0.004, t = 2.5, P = 0.026, df = 15. iii, 27.31 ± 1.34 pS, t = 0.63, P = 0.54, df = 15. Unpaired Student’s t-test compared with GluD2 in 100 µM dopamine. **d**, no ligand and no-protein controls. Example: 30-minute traces from bilayers in 10 mM dopamine, without the addition of GluD2 (*left,* n = 7) and from bilayers in the absence of ligand with GluD2 (*right*, n = 7). All repeats are independent bilayer recordings.

**Extended Data Fig. 2.**
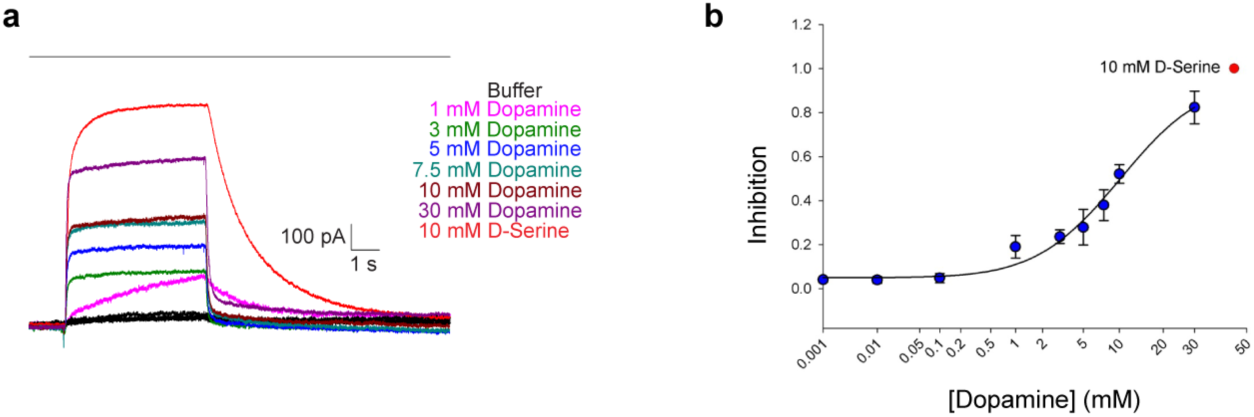
Dose-response of GluD2_Lc_ to dopamine. **a**, Representative whole-cell current traces recorded in response to multiple concentrations of dopamine The bar indicates the perfusion of the ligand. **b**, Dose-response curve for dopamine on GluD2_Lc_. The IC_50_ of dopamine against GluD2_Lc_ was measured to be 8.5 ± 1.97 mM. 10 nM (n = 6), 100 nM (n = 6), 1 µM (n = 6), 10 µM (n = 8), 0.1 mM (n = 8), 1 mM (n = 8), 3 mM (n = 7), 5 mM (n = 7), 7.5 mM (n = 7), 10 mM (n = 10), and 30 mM (n = 5). mean ± s.e.m. Inhibition is normalized to 10 mM D-serine. The holding potential for recordings was-60 mV.

**Extended Data Fig. 3.**
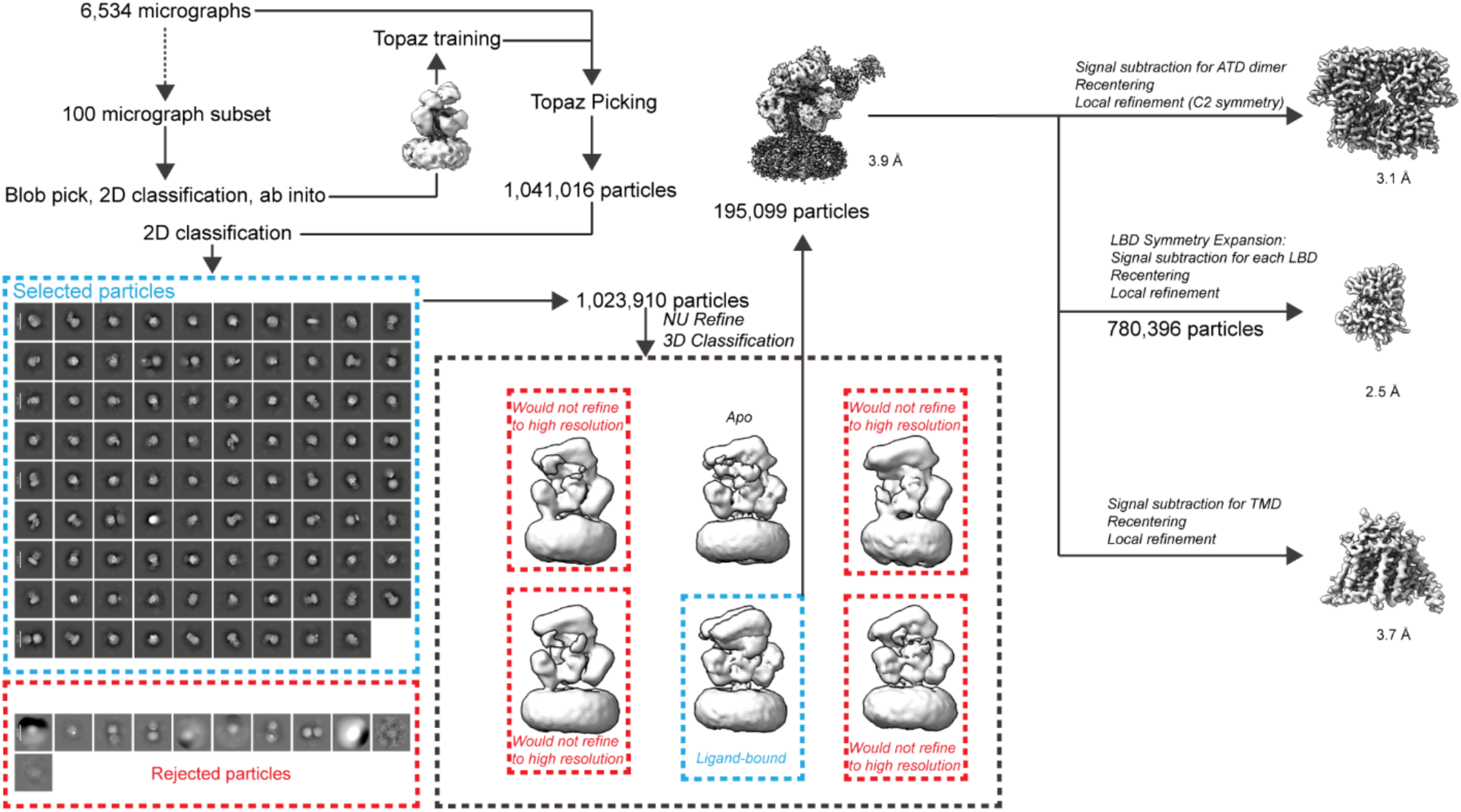
Cryo-EM image processing workflow for GluD2-dopamine. The Cryosparc workflow used to produce the final GluD2-dopamine maps.

**Extended Data Fig. 4.**
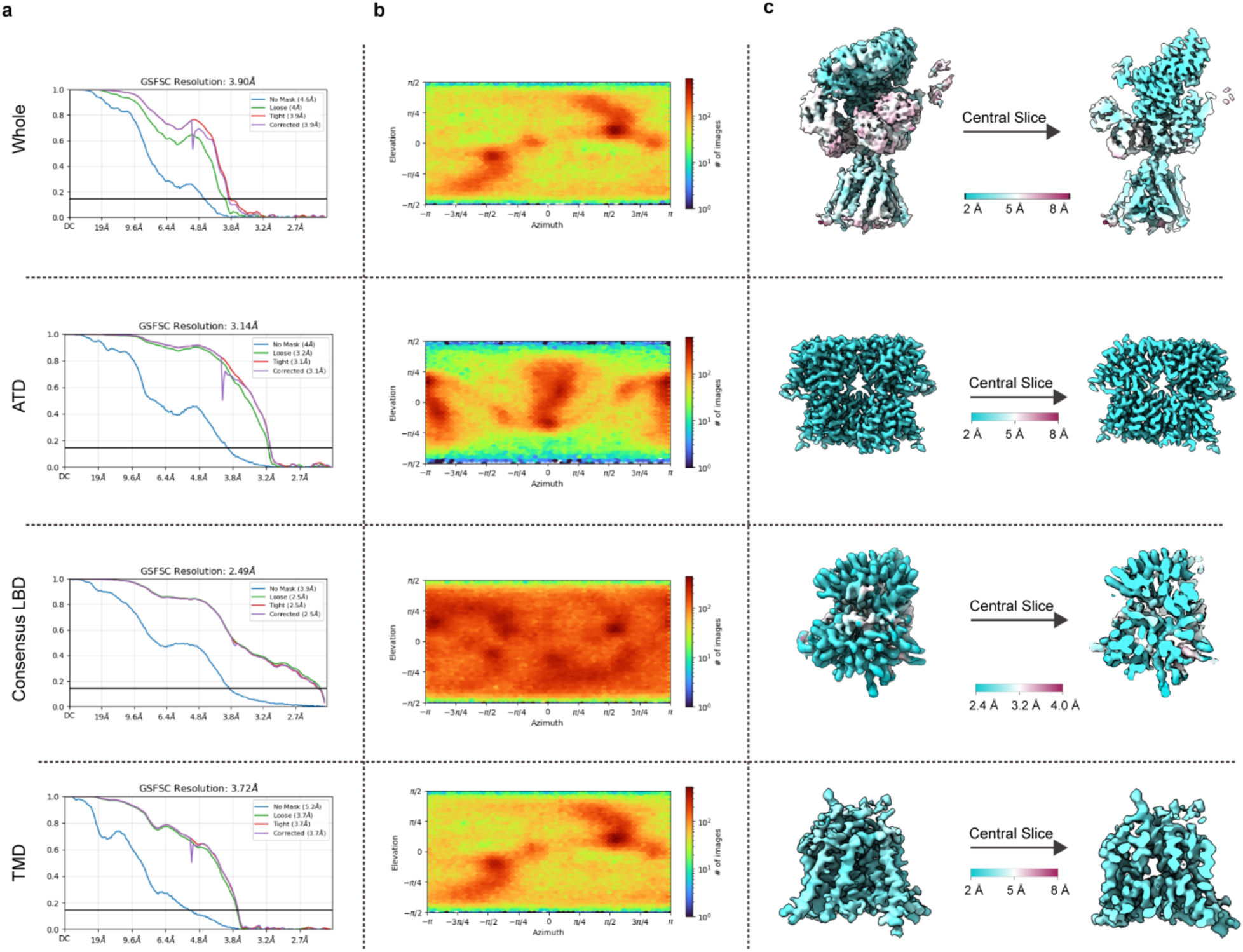
Overall map and local qualities for GluD2-dopamine. a, Gold standard Fourier shell correlation (GSFSC) curves for each whole and local map. The horizontal black line is the Fourier shell correlation (FSC) = 0.143. The y-axis is FSC; x is resolution in Å. **b,** Heat maps of the distribution of particle orientation for each map. **c,** Local resolution maps computed for each voxel, FSC = 0.143.

**Extended Data Fig. 5.**
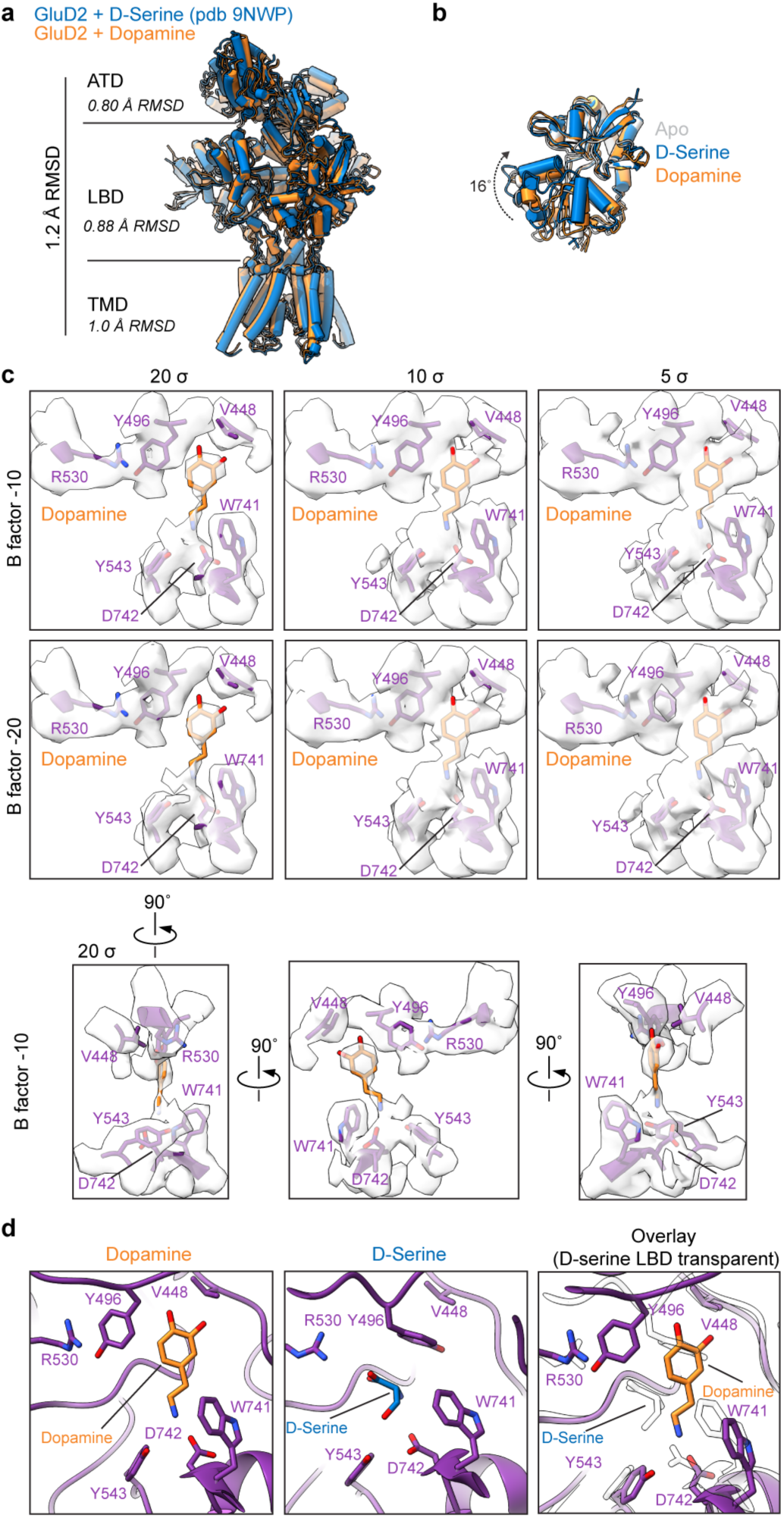
Comparison of GluD2 architecture and binding of D-serine and dopamine. a, Comparison of the overall and local architectures of GluD2-dopamine to GluD2-D-serine (PDB: 9NWP). RMSD, root mean square deviation. **b,** Side-view comparison of the LBD of Apo GluD2 (PDB: 9NWO), GluD2-dopamine and GluD2-D-serine. **c.** Model (purple) of the binding pose of dopamine (orange) with the associated local EM density map at multiple threshold values (σ, x-axis) and B factor levels (y-axis). **d.** Model of the binding pose of dopamine and the theoretical binding pose of D-serine (blue) within the LBD of GluD2-dopamine.

**Extended Data Fig. 6.**
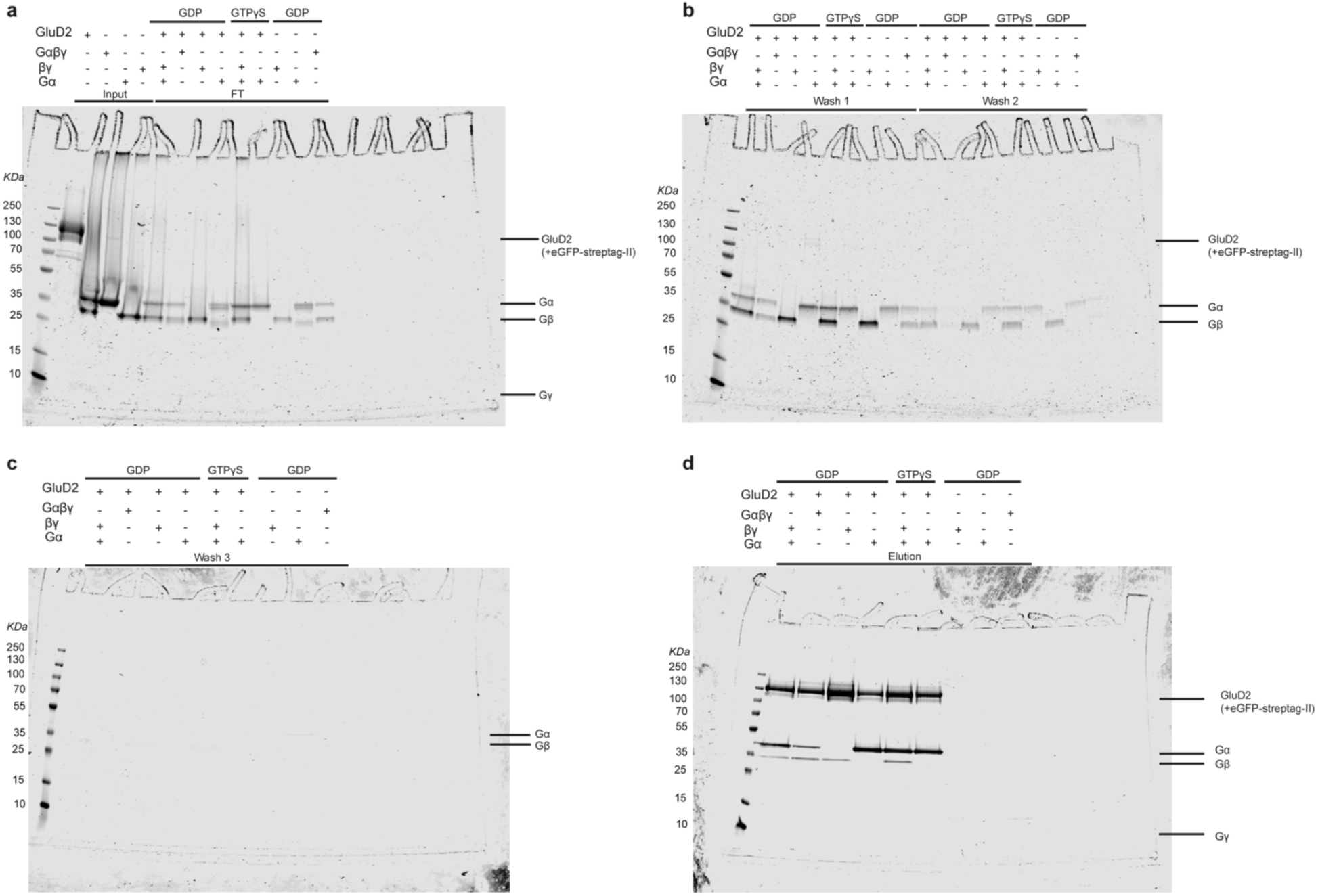
Example workflow for GluD2 G-protein complex reconstitutions. **a**, Input proteins and flowthrough (FT) for GluD2-eGFP-StrepTag II pull-downs. **b**, Washes 1 and 2. **c**, Wash 3. **d,** Elution of GluD2-eGFP-StrepTag II complexes. n = 3.

**Extended Data Fig. 7.**
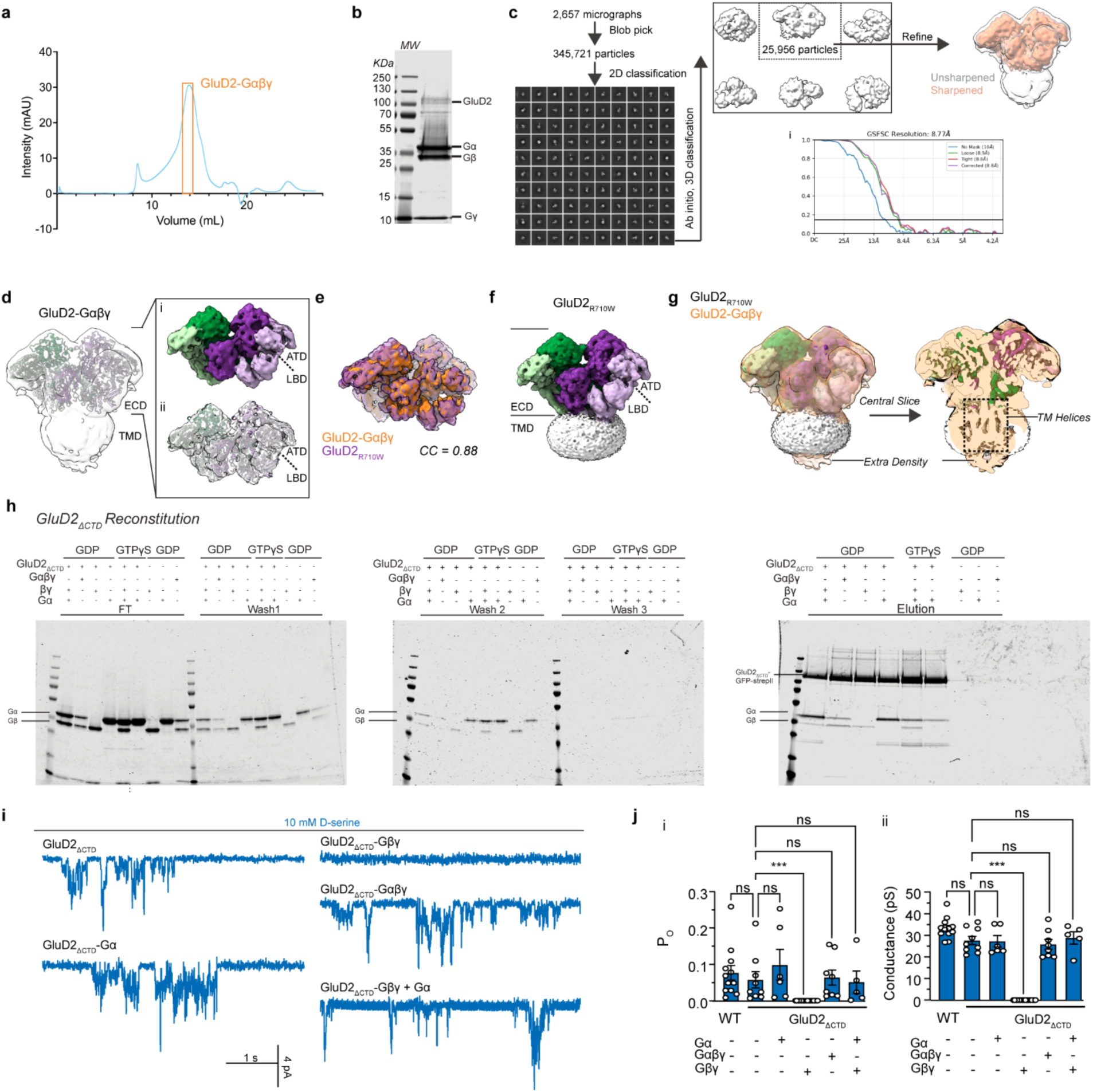
Mechanisms of modulation of GluD2 by G-proteins. **a**, Size-exclusion chromatograph of GluD2-Gαβγ. **b,** Example gel of purified GluD2-Gαβγ. **c,** The Cryosparc workflow used to produce the final GluD2-Gαβγ.maps with its corresponding GSFSC curve (i). **d**, GluD2-Gαβγ with the unsharpened composite map overlaid. Inset shows the surface representation (i) and the unsharpened local map (ii). **e**, Comparison of GluD2-Gαβγ (orange) with GluD2_R710W_ (purple, PDB: 9OOO) extracellular domains. **f**, Full-length map of GluD2_R710W_. **g**, Full-length map and central slice of GluD2_R710W_ with the unsharpened composite map of GluD2-Gαβγ. **h,** Input proteins and flowthrough (FT) for GluD2_ΔCTD_-eGFP-StrepTag II pull-downs (*left*), washes 1 and 2 (*middle*), wash and elution (*right*) of GluD2_ΔCTD_-eGFP-StrepTag II complexes (n = 3). **i,** Example current traces in the presence of 10 mM D-serine of GluD2-ΔCTD unbound by G-proteins (n = 9), bound to Gα (n *=* 6), Gβγ (n = 19), Gαβγ (n = 8), or Gα + Gβγ (n = 5). **j,** P_O_ summary (i), G-protein unbound WT GluD2 (0.070 ± 0.013), G-protein unbound ΔCTD: 0.059 ± 0.022, P > 0.99, mean rank difference =-5.1, Z = 0.66; ΔCTD-Gα: 0.10 ± 0.04, P > 0.99, mean rank difference =-4.8, Z = 0.49; ΔCTD-Gβγ: 0.00 ± 0.00, P = 0.0004, mean rank difference = 30, Z = 3.9; ΔCTD-Gαβγ: 0.065 ± 0.021, P > 0.99, mean rank difference =-3.2, Z = 0.35; ΔCTD-Gα+Gβγ: 0.053 ± 0.030, P > 0.99, mean rank difference = 3.1, Z = 0.29. Kruskal-Wallis test (Kruskal-Wallis statistic = 42, P < 0.00010) with Dunn’s correction relative to G-protein unbound GluD2_ΔCTD_. Conductance summary (ii), G-protein unbound WT GluD2 (25.44 ± 2.08 pS, n = 18), G-protein unbound ΔCTD: 27.64 ± 1.90 pS, P > 0.99, mean rank difference = 4.44, Z = 0.58; ΔCTD-Gα: 27.25 ± 2.58 pS, P > 0.99, mean rank difference = 1.33, Z = 0.14; ΔCTD-Gβγ: 0.00 ± 0.00 pS, P < 0.00010, mean rank difference = 35.00, Z = 4.6; ΔCTD-Gαβγ: 25.83 ± 2.25 pS, P > 0.99, mean rank difference = 4.5, Z = 0.50; ΔCTD-Gα+Gβγ: 28.80 ± 2.79 pS, P > 0.99, mean rank difference =-1.8, Z = 0.17. Kruskal-Wallis test with Dunn’s correction relative to G-protein unbound GluD2_ΔCTD_ (Kruskal-Wallis statistic = 41, P < 0.00010).

**Extended Data Fig. 8.**
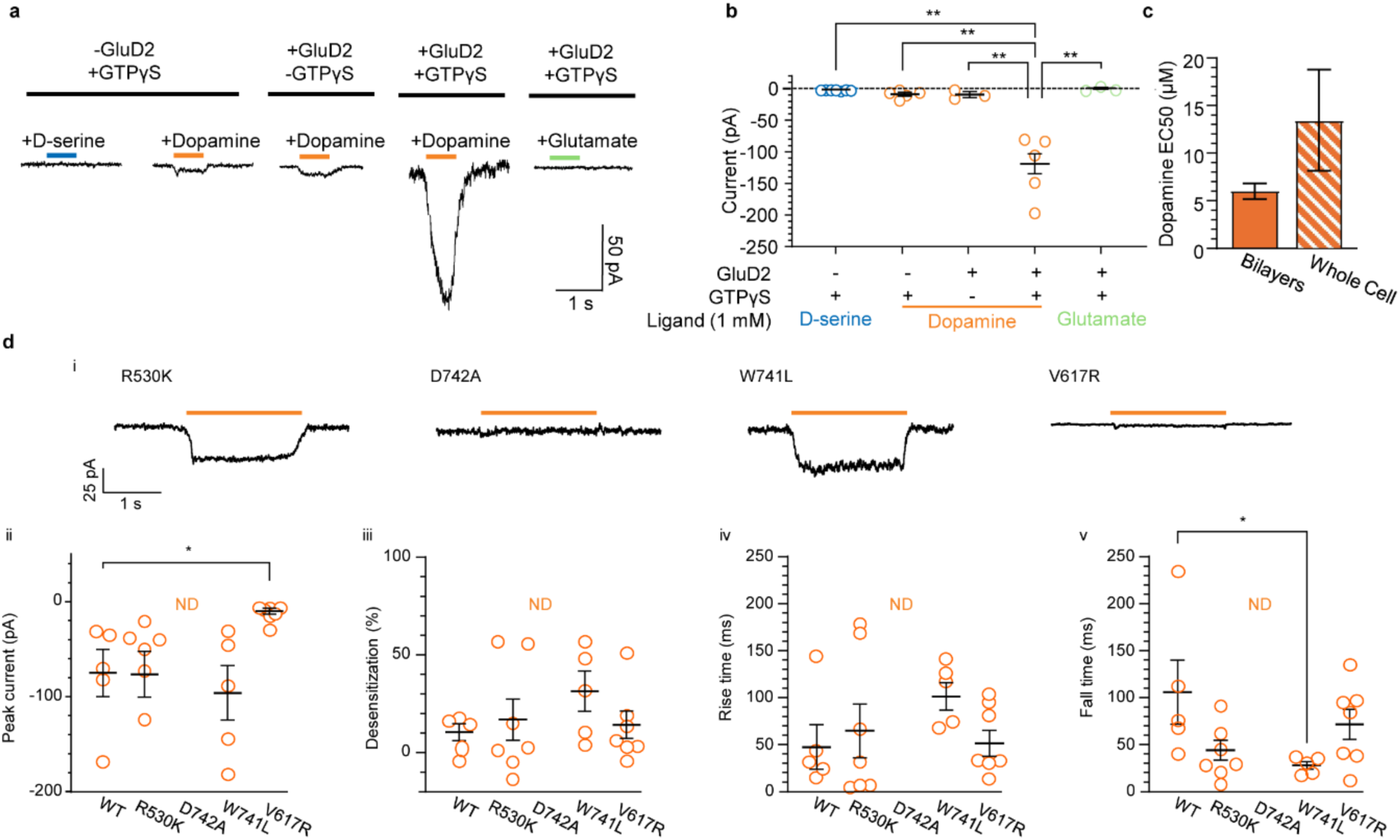
Additional whole-cell recordings. a,b, Representative whole-cell traces (a) and associated summary (b) of macroscopic GluD2 currents elicited by the ligands and conditions as shown. Uninfected, GTPγS (100 µM), D-serine (1 mM): n = 7, P = 0.0049, 95% CI of difference =-159.3 to-50.26, df = 4.0; Uninfected, GTPγS, dopamine (1 mM): n = 5, P = 0.0067, 95% CI of difference =-154.1 to-42.92, df = 4.3; GluD2, dopamine: n = 3, P = 0.0034, 95% CI of difference =-149.1 to-46.52, df = 4.6; GluD2, GTPγS, glutamate (1 mM): n = 3, P = 0.0048, 95% CI of difference =-161.9 to-51.63, df = 4.2. ANOVA (F = 51, P = 0.00030) with Dunnett’s correction relative to GluD2 + GTPγS + dopamine. **c**, Comparison of GluD2 EC_50_ values for dopamine derived from bilayer recordings vs whole-cell recordings (mean ± s.e.m.). t = 1.4, P = 0.17, df = 42. Unpaired Welch’s t-test. **d**, Representative whole-cell traces (i) and associated summaries of peak current amplitude (ii), desensitization (iii), rise time (iv) and fall time (v) of GluD2 mutations (R530K, n = 7; D742A, n = 8; W741L, n = 5; V617R, n = 6) compared to WT (n = 5). ND, not determined. ii, WT:-74.55 ± 24.69 pA; R530K:-76.79 ± 24.78 pA, P > 0.99, mean rank difference = 0.03, Z = 0.0069; W741:-95.42 ± 28.61 pA, P > 0.99, mean rank difference = 1.80, Z = 0.40; V617R:-9.69 ± 3.17, P = 0.02, mean rank difference =-11.26, Z = 2.7. Kruskal-Wallis test (Kruskal-Wallis statistic = 14, P = 0.0029) with Dunn’s correction relative to WT. iii, WT: 10.94 ± 4.32%; R530K, 16.81 ± 10.79%, P = 0.94, 95% CI of difference =-40.24 to 28.50, df = 7.8; W741L: 31.85 ± 10.29%, P = 0.28, 95% CI of difference =-58.84 to 17.03, df = 5.4, V617R: 14.67 ± 6.96%, P = 0.95, 95% CI of difference =-27.36 to 19.91, df = 9.4. ANOVA test (F = 0.99, P = 0.42) with Dunnett’s correction relative to WT. iv, WT: 47.46 ± 23.63 ms; R530K: 66.47 ± 28.93 ms, P = 0.94, 95% CI of difference =-124.70 to 86.65, df = 10; W741L: 101.1 ± 14.55 ms, P = 0.24, 95% CI of difference =-138.5 to 31.10, df = 6.7; V617R: 51.44 ± 13.74 ms, P = 0.99, 95% CI of difference =-87.50 to 79.53, df = 6.7. ANOVA test (F = 1.2, P = 0.36) with Dunnett’s correction relative to WT. v, WT: 99.50 ± 33.56 ms; R530K: 43.02 ± 10.72 ms, P = 0.25, mean rank difference = 7.14, Z = 1.7; W741L: 22.62 ± 3.90 ms, P = 0.019, mean rank difference = 12.20, Z = 2.7; V617R: 65.71 ± 15.94 ms, P > 0.99, mean rank difference = 3.00, Z = 0.72. Kruskal-Wallis test (Kruskal-Wallis statistic = 8.8, P = 0.033) with Dunn’s correction relative to WT. All repeats are from individual, separate cells.

**Extended Data Fig. 9.**
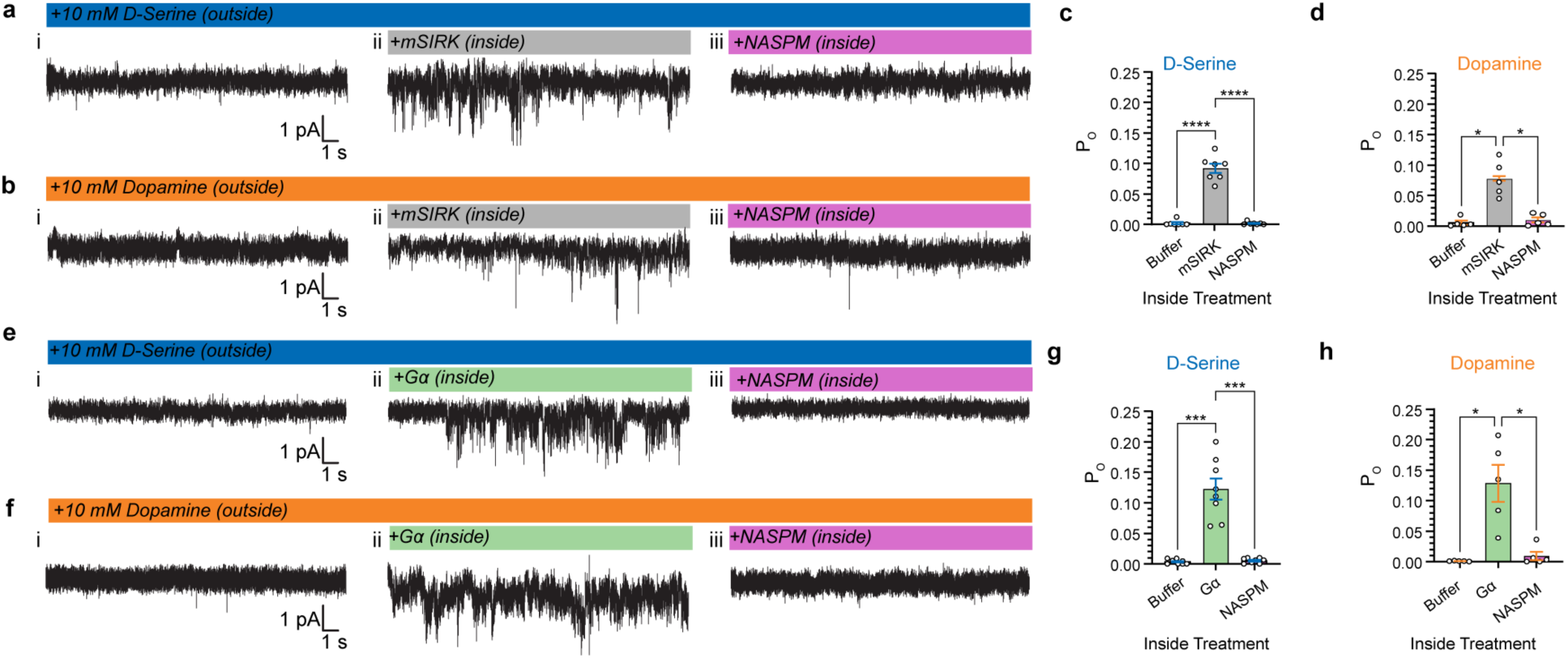
Inside-out recordings from GluD2. **a,b,** representative unitary current traces recorded from inside-out patches with D-serine (**a**) or dopamine (**b**) without (i), or with intracellular mSIRK (ii) or mSIRK + NASPM (iii)**. c,** Mean Po ± s.e.m. of baseline D-serine currents, baseline: 0.002 ± 0.002; intracellular mSIRK: 0.092 ± 0.008; intracellular mSIRK + NASPM: 0.002 ± 0.001, n = 7. Baseline vs mSIRK P < 0.00010, 95% CI of difference =-0.114 to-0.066, df = 6; Baseline vs mSIRK + NASPM P < 0.00010, 95% CI of difference = 0.065 to 0.115, df = 6. Repeated-measures ANOVA (F = 160, P < 0.00010) with Tukey’s correction. **d,** Mean Po ± s.e.m. of baseline dopamine currents, baseline: 0.005 ± 0.003; intracellular mSIRK: 0.077 ± 0.013; intracellular mSIRK + NASPM: 0.010 ± 0.004, n = 5. Baseline vs mSIRK P = 0.016, 95% CI of difference =-0.123 to-0.021, df = 4; Baseline vs mSIRK + NASPM P = 0.013, 95% CI of difference = 0.022 to 0.113, df = 4. Repeated-measures ANOVA (F = 26, P = 0.0061) with Tukey’s correction. **e,f,** representative unitary current traces recorded from inside-out patches with D-serine (**e**) or dopamine (**f**) without (i), or with intracellular Gα (ii) or Gα + NASPM (iii). **g,** Mean Po ± s.e.m. of baseline D-serine currents, baseline: 0.004 ± 0.001; intracellular Gα: 0.123 ± 0.017; intracellular Gα + NASPM: 0.005 ± 0.001, n = 8. Baseline vs Gα P = 0.0006, 95% CI of difference =-0.170 to-0.069, df = 8; Baseline vs Gα + NASPM P < 0.00010, 95% CI of difference = 0.069 to 0.166, df = 8. Repeated-measures ANOVA (F = 49, P = 0.00020) with Tukey’s correction. **h,** Mean P_o_ ± s.e.m. of baseline dopamine currents, baseline: 0.0007 ± 0.0003; intracellular Gα: 0.129 ± 0.031; intracellular Gα + NASPM: 0.009 ± 0.007, n = 5. Baseline vs Gα P = 0.030, 95% CI of difference =-0.236 to-0.019, df = 4; Baseline vs Gα + NASPM P = 0.027, 95% CI of difference = 0.020 to 0.218, df = 4. Repeated-measures ANOVA (F = 18, P = 0.011) with Tukey’s correction. All repeats are from individual, separate cells.

**Extended Data Fig. 10.**
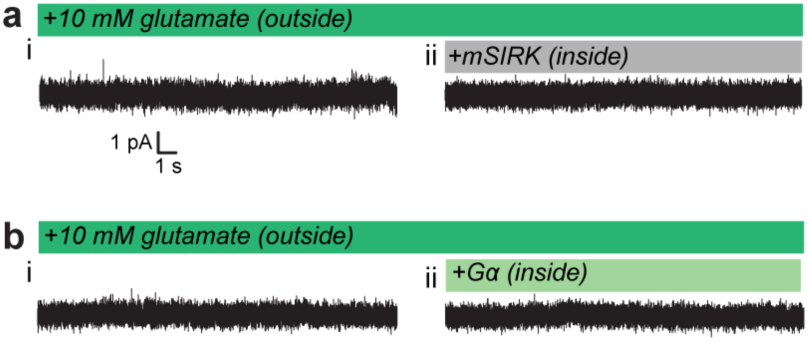
Controls for inside-out recordings. **a**, Representative trace of GluD2 exposed to 10 mM glutamate outside with no intracellular treatment (i) and then with intracellular mSIRK (ii); n = 5. **b**, Representative trace of GluD2 exposed to 10 mM glutamate outside with no intracellular treatment (i) and then with intracellular Gα (ii); n = 5. All repeats are from individual, separate cells.

---

**Extended Data Table 1.**
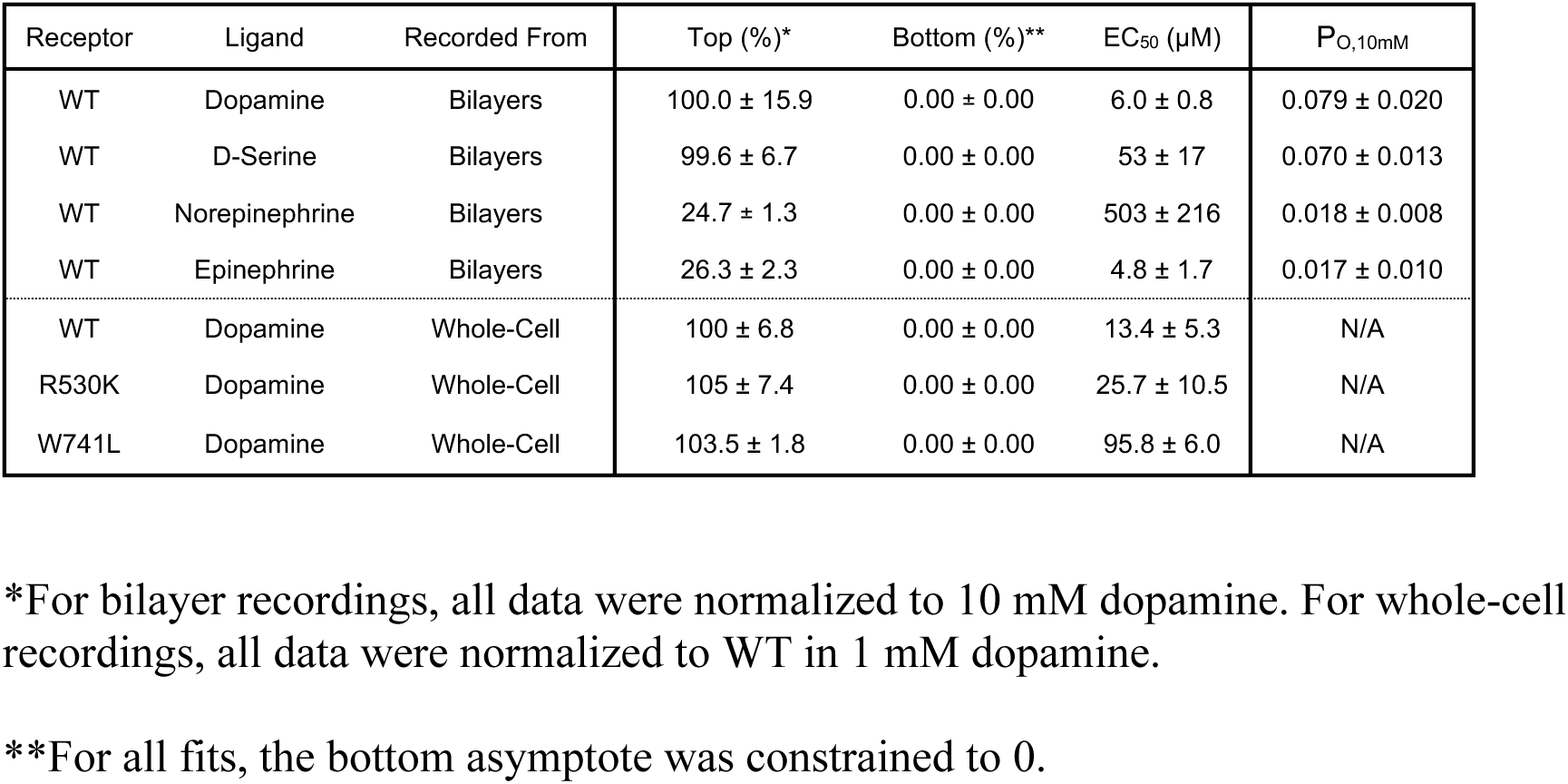
Electrophysiology dose-response statistics.

**Extended Data Table 2.**
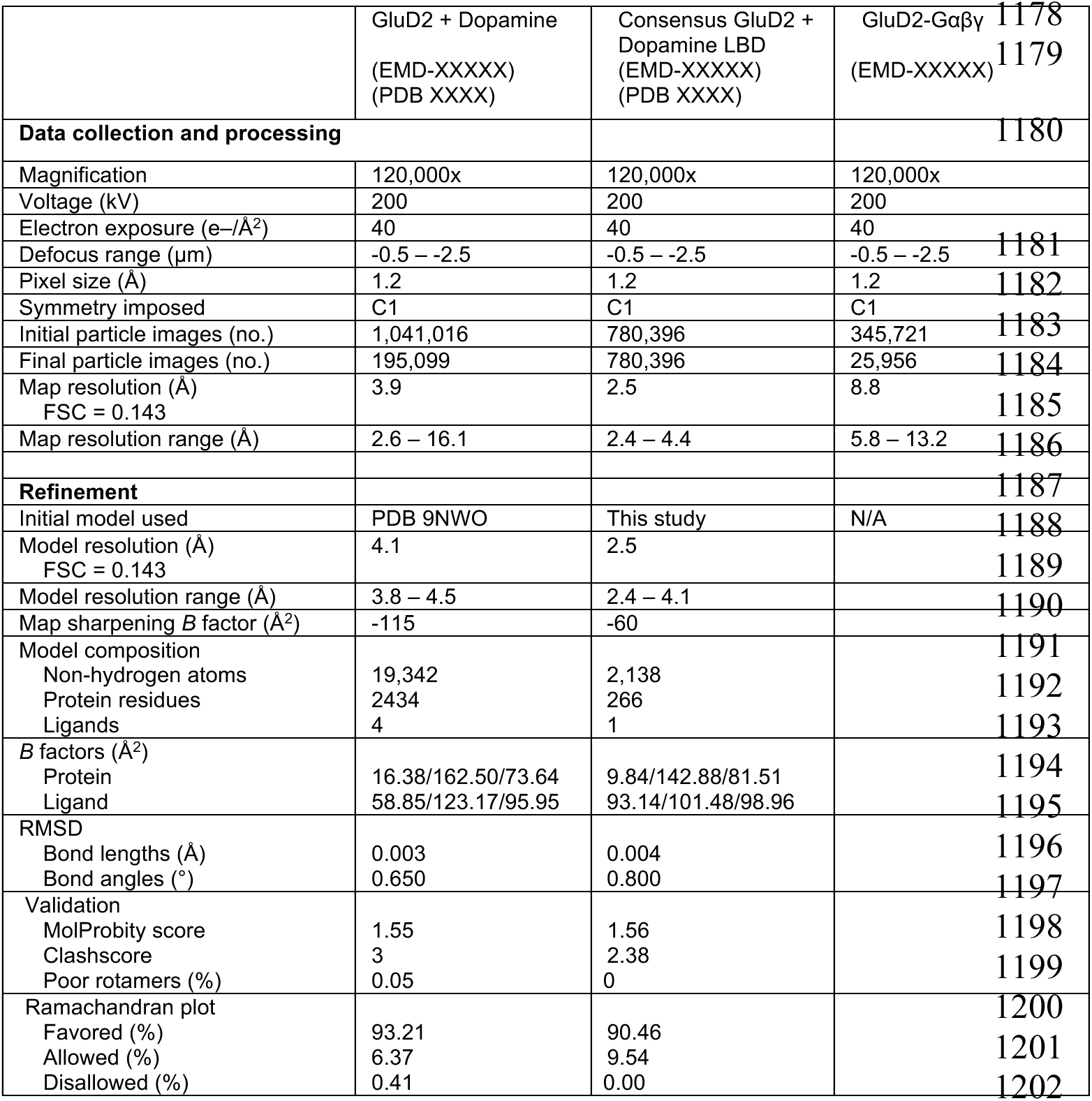
Cryo-EM data collection, refinement, and validation statistics.

|  | GluD2 + Dopamine<br>(EMD-XXXXX)<br>(PDB XXXX) | Consensus GluD2 + Dopamine LBD<br>(EMD-XXXXX)<br>(PDB XXXX) | GluD2-Gαβγ<br>(EMD-XXXXX) |
|---|---|---|---|
| **Data collection and processing** |  |  |  |
| Magnification | 120,000x | 120,000x | 120,000x |
| Voltage (kV) | 200 | 200 | 200 |
| Electron exposure (e–/Å<sup>2</sup>) | 40 | 40 | 40 |
| Defocus range (μm) | -0.5 – -2.5 | -0.5 – -2.5 | -0.5 – -2.5 |
| Pixel size (Å) | 1.2 | 1.2 | 1.2 |
| Symmetry imposed | C1 | C1 | C1 |
| Initial particle images (no.) | 1,041,016 | 780,396 | 345,721 |
| Final particle images (no.) | 195,099 | 780,396 | 25,956 |
| Map resolution (Å)<br>FSC = 0.143 | 3.9 | 2.5 | 8.8 |
| Map resolution range (Å) | 2.6 – 16.1 | 2.4 – 4.4 | 5.8 – 13.2 |
| **Refinement** |  |  |  |
| Initial model used | PDB 9NWO | This study | N/A |
| Model resolution (Å)<br>FSC = 0.143 | 4.1 | 2.5 |  |
| Model resolution range (Å) | 3.8 – 4.5 | 2.4 – 4.1 |  |
| Map sharpening *B* factor (Å<sup>2</sup>) | -115 | -60 |  |
| Model composition<br>Non-hydrogen atoms<br>Protein residues<br>Ligands | 19,342<br>2434<br>4 | 2,138<br>266<br>1 |  |
| *B* factors (Å<sup>2</sup>)<br>Protein<br>Ligand | 16.38/162.50/73.64<br>58.85/123.17/95.95 | 9.84/142.88/81.51<br>93.14/101.48/98.96 |  |
| RMSD<br>Bond lengths (Å)<br>Bond angles (°) | 0.003<br>0.650 | 0.004<br>0.800 |  |
| Validation<br>MolProbity score<br>Clashscore<br>Poor rotamers (%) | 1.55<br>3<br>0.05 | 1.56<br>2.38<br>0 |  |
| Ramachandran plot<br>Favored (%)<br>Allowed (%)<br>Disallowed (%) | 93.21<br>6.37<br>0.41 | 90.46<br>9.54<br>0.00 |  |

