## Extended Data for "GluDs are ionotropic dopamine receptors tuned by G-proteins"

### Extended Data Figures

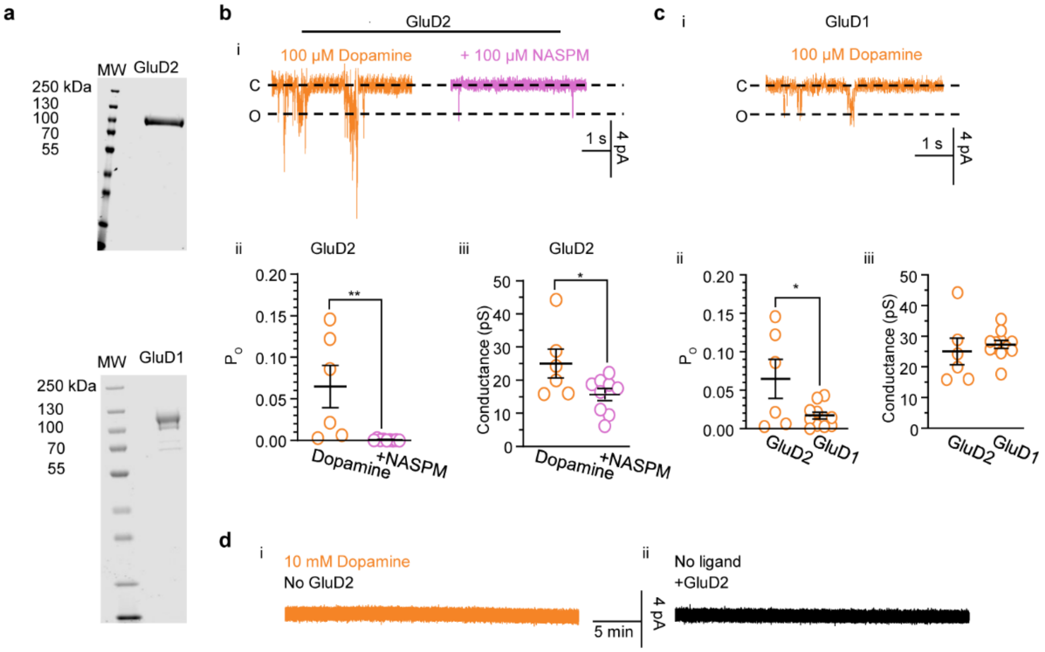

**Extended Data Fig. 1 | Comparison of dopaminergic activation of isolated GluD1 and GluD2.** **a,** Example gels of purified GluD2 *(top)* and GluD1 *(bottom)*. **b,** Example current traces (i) of GluD2 in response to 100 µM dopamine without (n = 6) or with (n = 9) the addition of 100 µM NASPM, with associated summaries (mean ± s.e.m.) of $P_O$ (ii) and conductance (iii). ii, 100 µM dopamine: 0.065 ± 0.025; 100 µM dopamine + 100 µM NASPM: 0.00070 + 0.00030, t = 3.1, P = 0.0079, df = 13. iii, 100 µM dopamine: 25.02 + 4.37 pS, t = 2.2, P = 0.048, df = 13. Unpaired Student's t-test. **c,** Example current traces (i) of GluD1 in response to 100 µM dopamine (n = 11) with associated summaries (mean ± s.e.m.) of $P_O$ (ii) and conductance (iii). ii, 0.017 ± 0.004, t = 2.5, P = 0.026, df = 15. iii, 27.31 ± 1.34 pS, t = 0.63, P = 0.54, df = 15. Unpaired Student's t-test compared with GluD2 in 100 µM dopamine. **d**, no ligand and no-protein controls. Example: 30-minute traces from bilayers in 10 mM dopamine, without the addition of GluD2 (*left,* n = 7) and from bilayers in the absence of ligand with GluD2 (*right*, n = 7). All repeats are independent bilayer recordings.

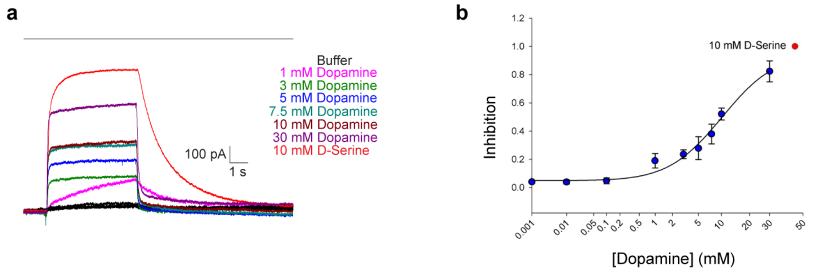

**Extended Data Fig. 2 | Dose-response of $GluD2_{Lc}$ to dopamine. a**, Representative whole-cell current traces recorded in response to multiple concentrations of dopamine The bar indicates the perfusion of the ligand. **b**, Dose-response curve for dopamine on $GluD2_{Lc}$. The $IC_{50}$ of dopamine against $GluD2_{Lc}$ was measured to be 8.5 ± 1.97 mM. 10 nM (n = 6), 100 nM (n = 6), 1 µM (n = 6), 10 µM (n = 8), 0.1 mM (n = 8), 1 mM (n = 8), 3 mM (n = 7), 5 mM (n = 7), 7.5 mM (n = 7), 10 mM (n = 10), and 30 mM (n = 5). mean ± s.e.m. Inhibition is normalized to 10 mM D-serine. The holding potential for recordings was -60 mV.

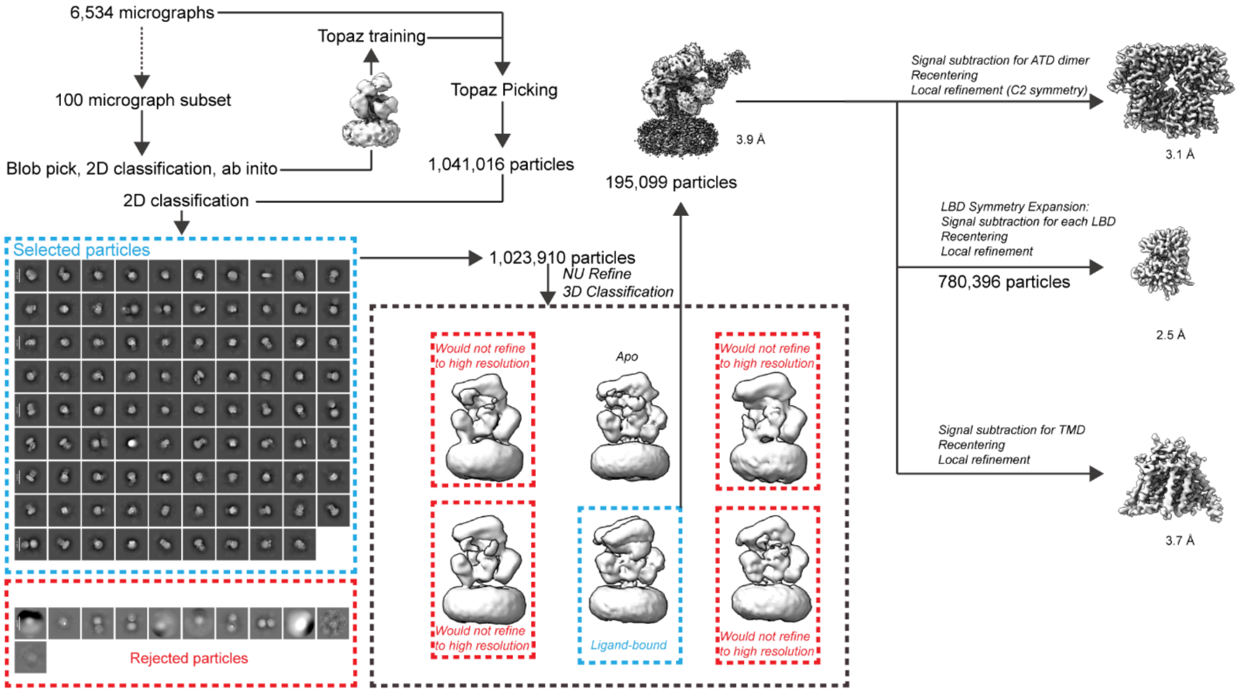

**Extended Data Fig. 3 | Cryo-EM image processing workflow for GluD2-dopamine.** The Cryosparc workflow used to produce the final GluD2-dopamine maps.

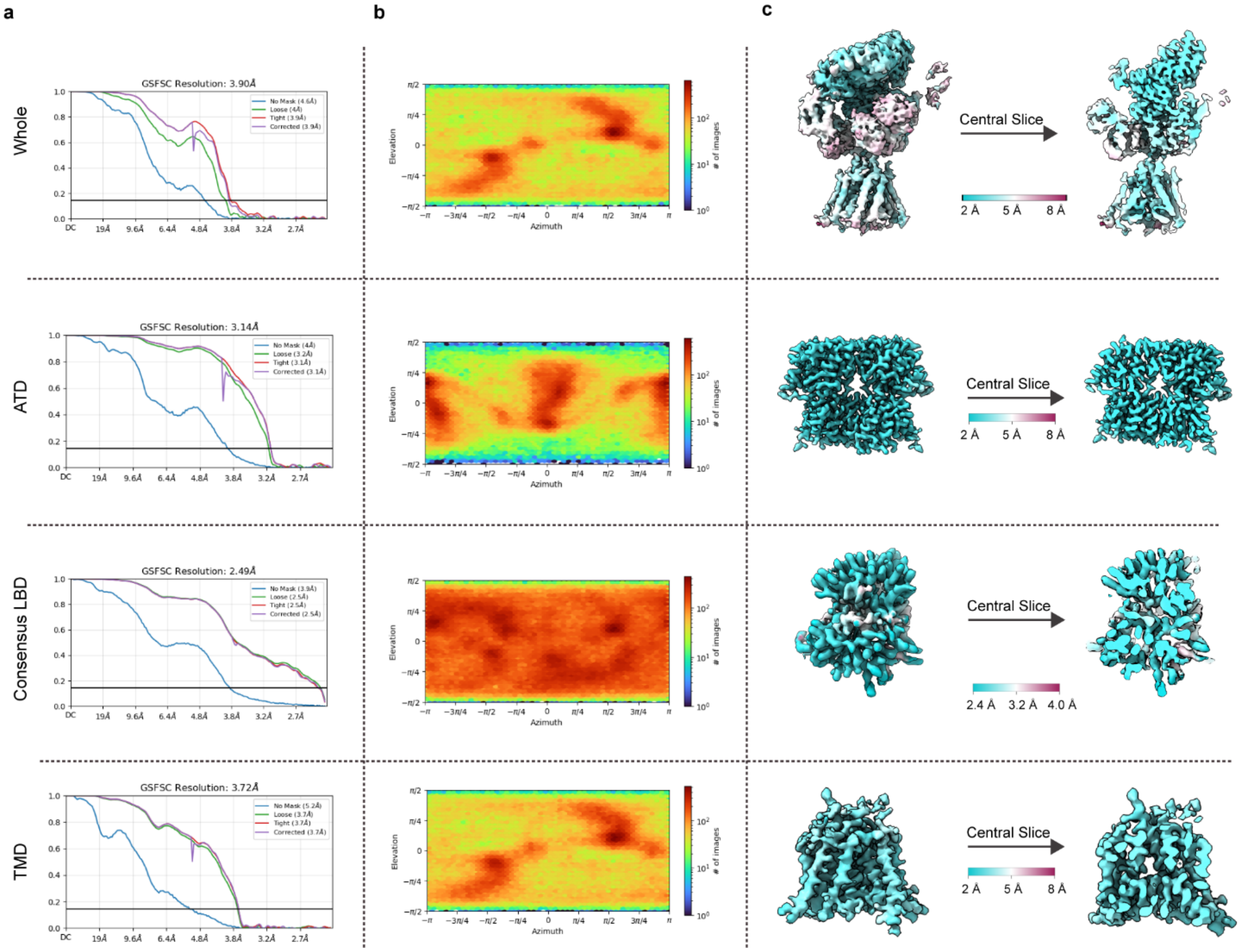

**Extended Data Fig. 4 | Overall map and local qualities for GluD2-dopamine. a,** Gold standard Fourier shell correlation (GSFSC) curves for each whole and local map. The horizontal black line is the Fourier shell correlation (FSC) = 0.143. The y-axis is FSC; x is resolution in Å. **b,** Heat maps of the distribution of particle orientation for each map. **c,** Local resolution maps computed for each voxel, FSC = 0.143.

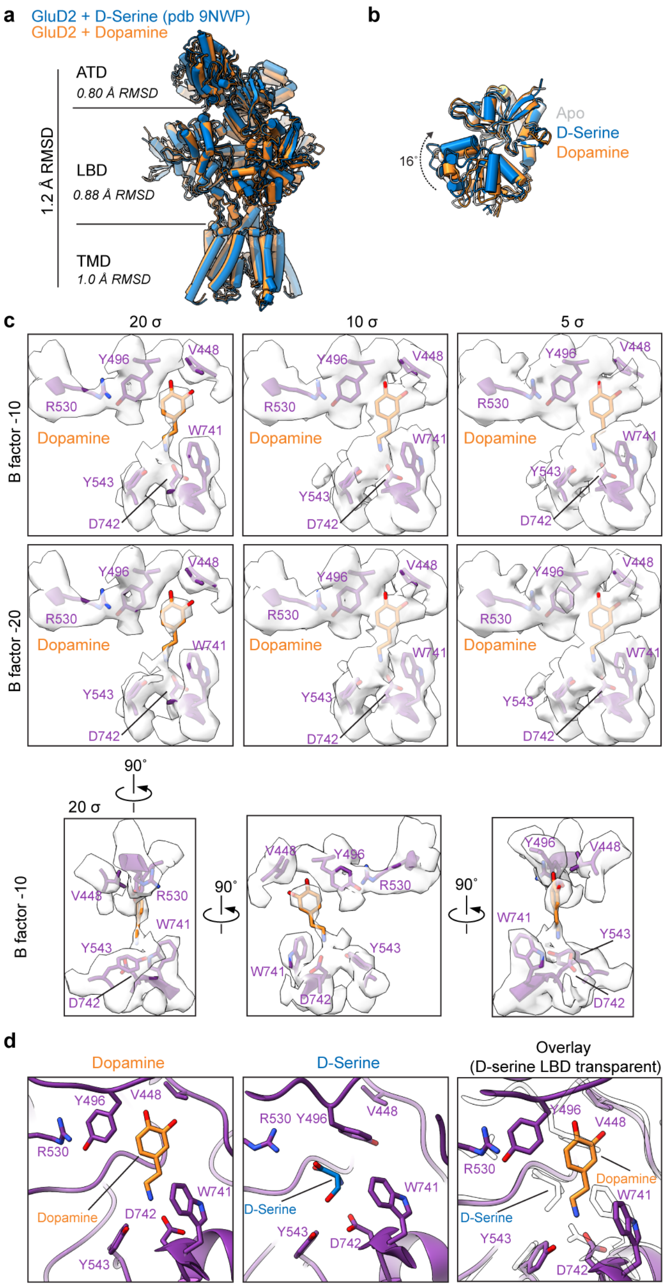

**Extended Data Fig. 5 | Comparison of GluD2 architecture and binding of D-serine and dopamine. a,** Comparison of the overall and local architectures of GluD2-dopamine to GluD2-D-serine (PDB: 9NWP). RMSD, root mean square deviation. **b,** Side-view comparison of the LBD of Apo GluD2 (PDB: 9NWO), GluD2-dopamine and GluD2-D-serine. **c.** Model (purple) of the binding pose of dopamine (orange) with the associated local EM density map at multiple threshold values (σ, x-axis) and B factor levels (y-axis). **d.** Model of the binding pose of dopamine and the theoretical binding pose of D-serine (blue) within the LBD of GluD2-dopamine.

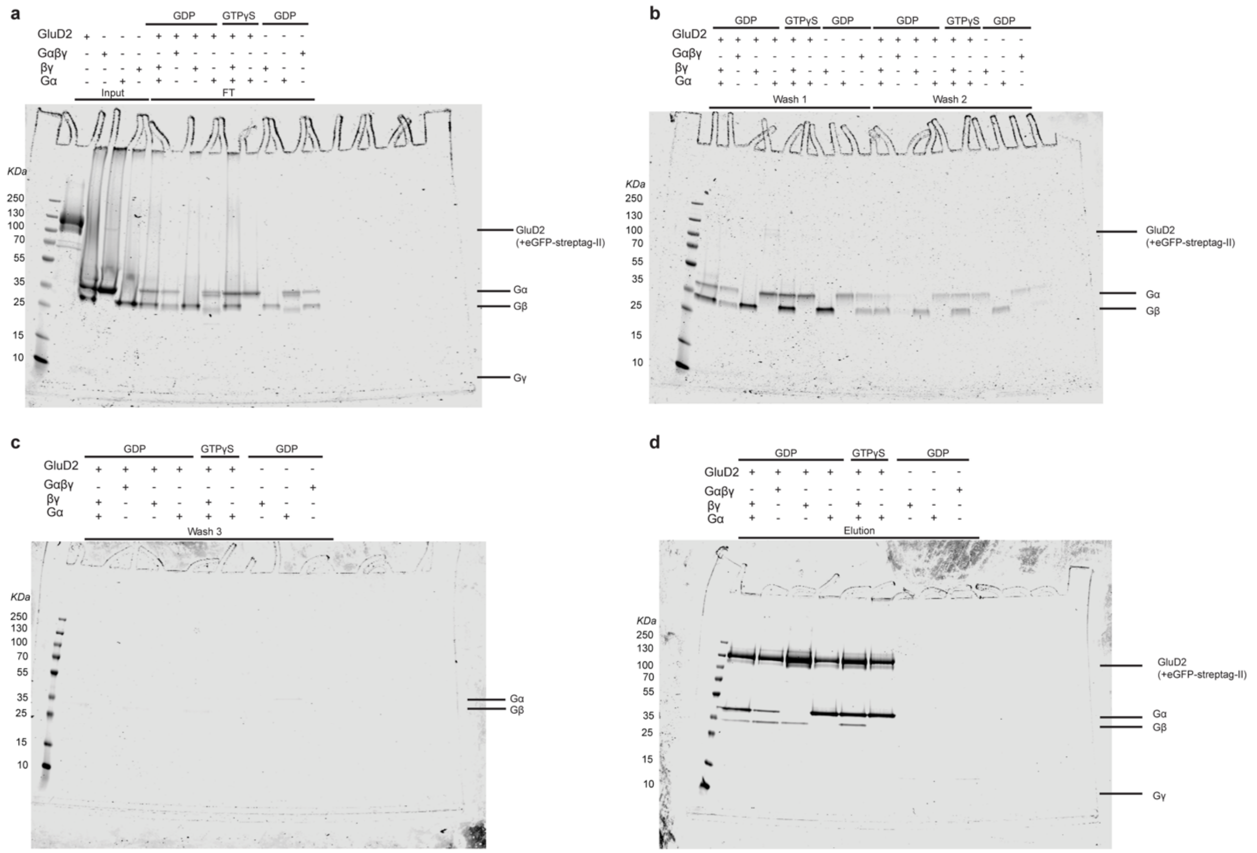

**Extended Data Fig. 6 | Example workflow for GluD2 G-protein complex reconstitutions. a**, Input proteins and flowthrough (FT) for GluD2-eGFP-StrepTag II pull-downs. **b**, Washes 1 and 2. **c**, Wash 3. **d,** Elution of GluD2-eGFP-StrepTag II complexes. n = 3.

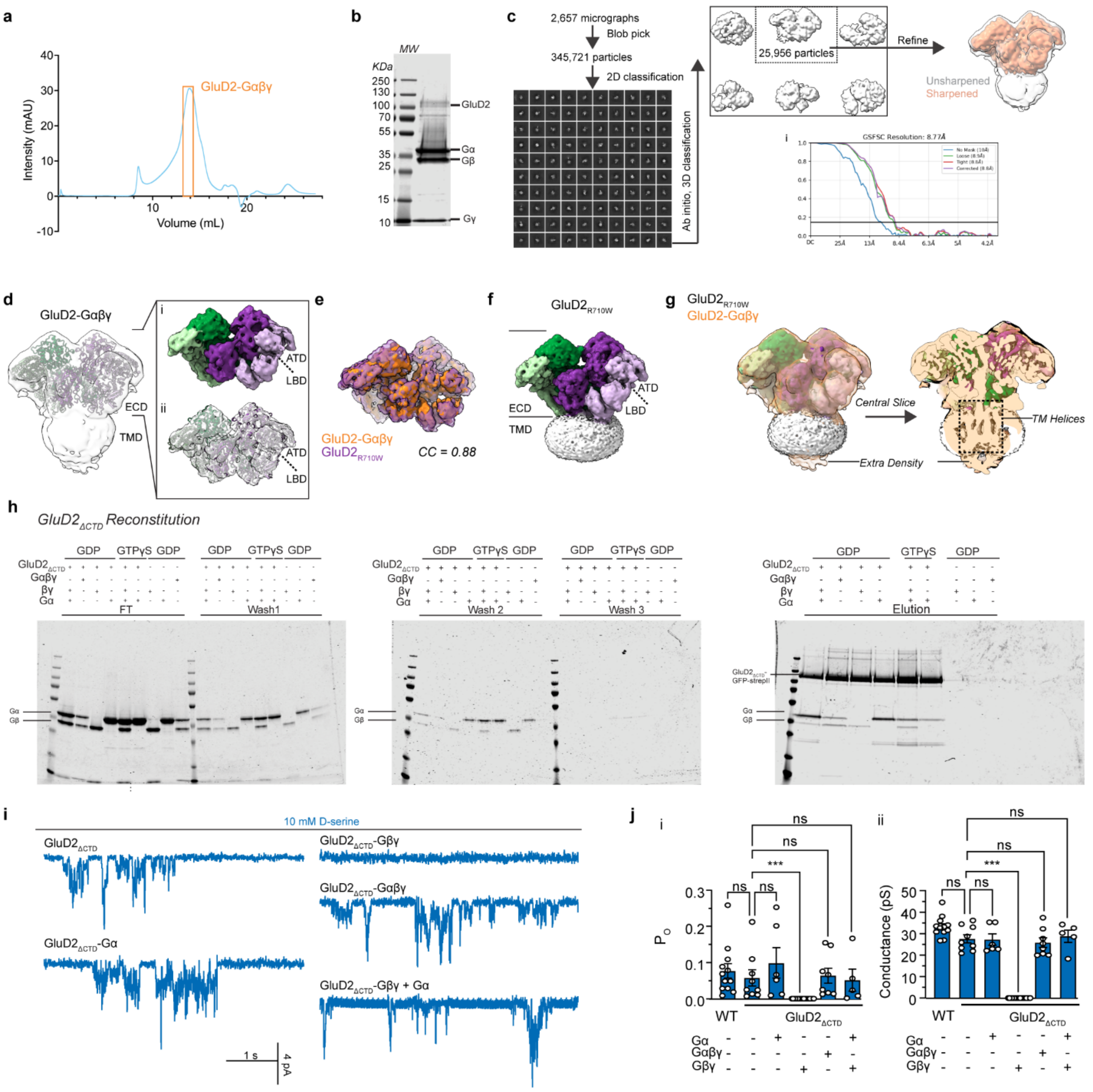

**Extended Data Fig. 7 | Mechanisms of modulation of GluD2 by G-proteins. a**, Size-exclusion chromatograph of GluD2-Gαβγ. **b,** Example gel of purified GluD2-Gαβγ. **c,** The Cryosparc workflow used to produce the final GluD2-Gαβγ.maps with its corresponding GSFSC curve (i). **d**, GluD2-Gαβγ with the unsharpened composite map overlaid. Inset shows the surface representation (i) and the unsharpened local map (ii). **e**, Comparison of GluD2-Gαβγ (orange) with GluD2$_{R710W}$ (purple, PDB: 9OOO) extracellular domains. **f**, Full-length map of GluD2$_{R710W}$. **g**, Full-length map and central slice of GluD2$_{R710W}$ with the unsharpened composite map of GluD2-Gαβγ. **h,** Input proteins and flowthrough (FT) for GluD2$_{\Delta CTD}$-eGFP-StrepTag II pull-downs (*left*), washes 1 and 2 (*middle*), wash and elution (*right*) of GluD2$_{\Delta CTD}$-eGFP-StrepTag II complexes (n = 3). **i,** Example current traces in the presence of 10 mM D-serine of GluD2-ΔCTD unbound by G-proteins (n = 9), bound to Gα (n = 6), Gβγ (n = 19), Gαβγ (n = 8), or Gα + Gβγ (n = 5). **j,** $P_O$ summary (i), G-protein unbound WT GluD2 (0.070 ± 0.013), G-protein unbound ΔCTD: 0.059 ± 0.022, P > 0.99, mean rank difference = -5.1, Z = 0.66; ΔCTD-Gα: 0.10 ± 0.04, P > 0.99, mean rank difference = -4.8, Z = 0.49; ΔCTD-Gβγ: 0.00 ± 0.00, P = 0.0004, mean rank difference = 30, Z = 3.9; ΔCTD-Gαβγ: 0.065 ± 0.021, P > 0.99, mean rank difference = -3.2, Z = 0.35; ΔCTD-Gα+Gβγ: 0.053 ± 0.030, P > 0.99, mean rank difference = 3.1, Z = 0.29. Kruskal-Wallis test (Kruskal-Wallis statistic = 42, P < 0.00010) with Dunn's correction relative to G-protein unbound GluD2$_{\Delta CTD}$. Conductance summary (ii), G-protein unbound WT GluD2 (25.44 ± 2.08 pS, n = 18), G-protein unbound ΔCTD: 27.64 ± 1.90 pS, P > 0.99, mean rank difference = 4.44, Z = 0.58; ΔCTD-Gα: 27.25 ± 2.58 pS, P > 0.99, mean rank difference = 1.33, Z = 0.14; ΔCTD-Gβγ: 0.00 ± 0.00 pS, P < 0.00010, mean rank difference = 35.00, Z = 4.6; ΔCTD-Gαβγ: 25.83 ± 2.25 pS, P > 0.99, mean rank difference = 4.5, Z = 0.50; ΔCTD-Gα+Gβγ: 28.80 ± 2.79 pS, P > 0.99, mean rank difference = -1.8, Z = 0.17. Kruskal-Wallis test with Dunn's correction relative to G-protein unbound GluD2$_{\Delta CTD}$ (Kruskal-Wallis statistic = 41, P < 0.00010).

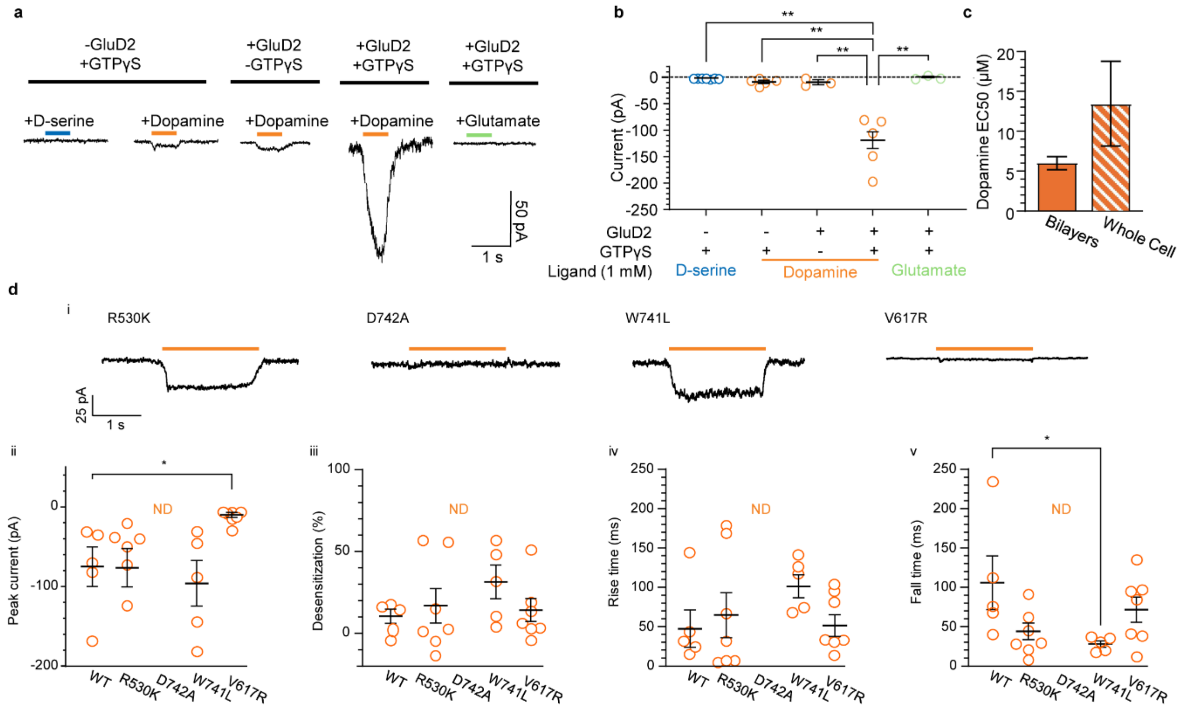

**Extended Data Fig. 8 | Additional whole-cell recordings. a,b,** Representative whole-cell traces (a) and associated summary (b) of macroscopic GluD2 currents elicited by the ligands and conditions as shown. Uninfected, GTPγS (100 μM), D-serine (1 mM): n = 7, P = 0.0049, 95% CI of difference = -159.3 to -50.26, df = 4.0; Uninfected, GTPγS, dopamine (1 mM): n = 5, P = 0.0067, 95% CI of difference = -154.1 to -42.92, df = 4.3; GluD2, dopamine: n = 3, P = 0.0034, 95% CI of difference = -149.1 to -46.52, df = 4.6; GluD2, GTPγS, glutamate (1 mM): n = 3, P = 0.0048, 95% CI of difference = -161.9 to -51.63, df = 4.2. ANOVA (F = 51, P = 0.00030) with Dunnett's correction relative to GluD2 + GTPγS + dopamine. **c**, Comparison of GluD2 $EC_{50}$ values for dopamine derived from bilayer recordings vs whole-cell recordings (mean ± s.e.m.). t = 1.4, P = 0.17, df = 42. Unpaired Welch's t-test. **d**, Representative whole-cell traces (i) and associated summaries of peak current amplitude (ii), desensitization (iii), rise time (iv) and fall time (v) of GluD2 mutations (R530K, n = 7; D742A, n = 8; W741L, n = 5; V617R, n = 6) compared to WT (n = 5). ND, not determined. ii, WT: -74.55 ± 24.69 pA; R530K: -76.79 ± 24.78 pA, P > 0.99, mean rank difference = 0.03, Z = 0.0069; W741: -95.42 ± 28.61 pA, $P > 0.99$, mean rank difference = 1.80, Z = 0.40; V617R: -9.69 ± 3.17, P = 0.02, mean rank difference = -11.26, Z = 2.7. Kruskal-Wallis test (Kruskal-Wallis statistic = 14, P = 0.0029) with Dunn's correction relative to WT. iii, WT: 10.94 ± 4.32%; R530K, 16.81 ± 10.79%, P = 0.94, 95% CI of difference = -40.24 to 28.50, df = 7.8; W741L: 31.85 ± 10.29%, P = 0.28, 95% CI of difference = -58.84 to 17.03, df = 5.4, V617R: 14.67 ± 6.96%, P = 0.95, 95% CI of difference = -27.36 to 19.91, df = 9.4. ANOVA test (F = 0.99, P = 0.42) with Dunnett's correction relative to WT. iv, WT: 47.46 ± 23.63 ms; R530K: 66.47 ± 28.93 ms, P = 0.94, 95% CI of difference = -124.70 to 86.65, df = 10; W741L: 101.1 ± 14.55 ms, P = 0.24, 95% CI of difference = -138.5 to 31.10, df = 6.7; V617R: 51.44 ± 13.74 ms, P = 0.99, 95% CI of difference = -87.50 to 79.53, df = 6.7. ANOVA test (F = 1.2, P = 0.36) with Dunnett's correction relative to WT. v, WT: 99.50 ± 33.56 ms; R530K: 43.02 ± 10.72 ms, P = 0.25, mean rank difference = 7.14, Z = 1.7; W741L: 22.62 ± 3.90 ms, P = 0.019, mean rank difference = 12.20, Z = 2.7; V617R: 65.71 ± 15.94 ms, $P > 0.99$, mean rank difference = 3.00, Z = 0.72. Kruskal-Wallis test (Kruskal-Wallis statistic = 8.8, P = 0.033) with Dunn's correction relative to WT. All repeats are from individual, separate cells.

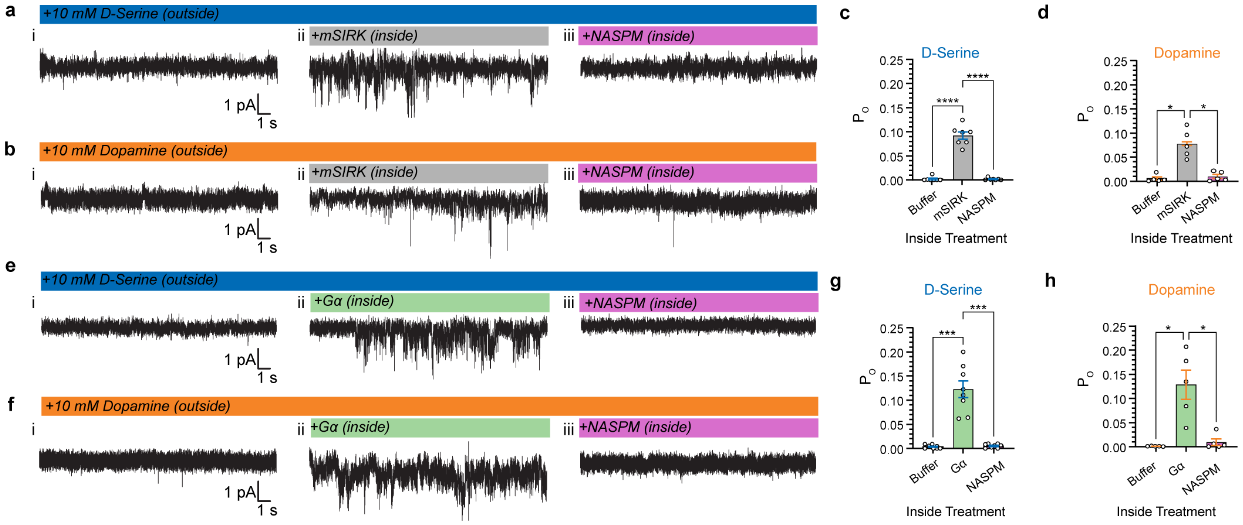

**Extended Data Fig. 9 | Inside-out recordings from GluD2. a,b,** representative unitary current traces recorded from inside-out patches with D-serine (**a**) or dopamine (**b**) without (i), or with intracellular mSIRK (ii) or mSIRK + NASPM (iii)**. c,** Mean Po ± s.e.m. of baseline D-serine currents, baseline: 0.002 ± 0.002; intracellular mSIRK: 0.092 ± 0.008; intracellular mSIRK + NASPM: 0.002 ± 0.001, n = 7. Baseline vs mSIRK P < 0.00010, 95% CI of difference = -0.114 to -0.066, df = 6; Baseline vs mSIRK + NASPM P < 0.00010, 95% CI of difference = 0.065 to 0.115, df = 6. Repeated-measures ANOVA (F = 160, P < 0.00010) with Tukey's correction. **d,** Mean Po ± s.e.m. of baseline dopamine currents, baseline: 0.005 ± 0.003; intracellular mSIRK: 0.077 ± 0.013; intracellular mSIRK + NASPM: 0.010 ± 0.004, n = 5. Baseline vs mSIRK P = 0.016, 95% CI of difference = -0.123 to -0.021, df = 4; Baseline vs mSIRK + NASPM P = 0.013, 95% CI of difference = 0.022 to 0.113, df = 4. Repeated-measures ANOVA (F = 26, P = 0.0061) with Tukey's correction. **e,f,** representative unitary current traces recorded from inside-out patches with D-serine (**e**) or dopamine (**f**) without (i), or with intracellular Gα (ii) or Gα + NASPM (iii). **g,** Mean Po ± s.e.m. of baseline D-serine currents, baseline: 0.004 ± 0.001; intracellular Gα: 0.123 ± 0.017; intracellular Gα + NASPM: 0.005 ± 0.001, n = 8. Baseline vs Gα P = 0.0006, 95% CI of difference = -0.170 to -0.069, df = 8; Baseline vs Gα + NASPM P < 0.00010, 95% CI of difference = 0.069

to 0.166, df = 8**.** Repeated-measures ANOVA (F = 49, P = 0.00020) with Tukey's correction. **h,** Mean $P_o$ ± s.e.m. of baseline dopamine currents, baseline: 0.0007 ± 0.0003; intracellular Gα: 0.129 ± 0.031; intracellular Gα + NASPM: 0.009 ± 0.007**,** n = 5. Baseline vs Gα P = 0.030, 95% CI of difference = -0.236 to -0.019, df = 4; Baseline vs Gα + NASPM P = 0.027, 95% CI of difference = 0.020 to 0.218, df = 4. Repeated-measures ANOVA (F = 18, P = 0.011) with Tukey's correction. All repeats are from individual, separate cells.

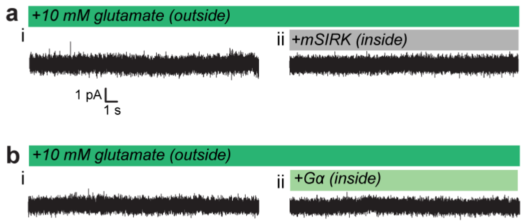

**Extended Data Fig. 10 | Controls for inside-out recordings. a**, Representative trace of GluD2 exposed to 10 mM glutamate outside with no intracellular treatment (i) and then with intracellular mSIRK (ii); n = 5. **b**, Representative trace of GluD2 exposed to 10 mM glutamate outside with no intracellular treatment (i) and then with intracellular Gα (ii); n = 5. All repeats are from individual, separate cells.

**Extended Data Table 1 | Electrophysiology dose-response statistics**

| Receptor | Ligand | Recorded From | Top (%)* | Bottom (%)** | $EC_{50}$ (μM) | $P_{O,10mM}$ |
|---|---|---|---|---|---|---|
| WT | Dopamine | Bilayers | 100.0 ± 15.9 | 0.00 ± 0.00 | 6.0 ± 0.8 | 0.079 ± 0.020 |
| WT | D-Serine | Bilayers | 99.6 ± 6.7 | 0.00 ± 0.00 | 53 ± 17 | 0.070 ± 0.013 |
| WT | Norepinephrine | Bilayers | 24.7 ± 1.3 | 0.00 ± 0.00 | 503 ± 216 | 0.018 ± 0.008 |
| WT | Epinephrine | Bilayers | 26.3 ± 2.3 | 0.00 ± 0.00 | 4.8 ± 1.7 | 0.017 ± 0.010 |
| WT | Dopamine | Whole-Cell | 100 ± 6.8 | 0.00 ± 0.00 | 13.4 ± 5.3 | N/A |
| R530K | Dopamine | Whole-Cell | 105 ± 7.4 | 0.00 ± 0.00 | 25.7 ± 10.5 | N/A |
| W741L | Dopamine | Whole-Cell | 103.5 ± 1.8 | 0.00 ± 0.00 | 95.8 ± 6.0 | N/A |

*For bilayer recordings, all data were normalized to 10 mM dopamine. For whole-cell recordings, all data were normalized to WT in 1 mM dopamine.

**For all fits, the bottom asymptote was constrained to 0.

**Extended Data Table 2 | Cryo-EM data collection, refinement, and validation statistics**

| | GluD2 + Dopamine (EMD-XXXXX) (PDB XXXX) | Consensus GluD2 + Dopamine LBD (EMD-XXXXX) (PDB XXXX) | GluD2-Gαβγ (EMD-XXXXX) |
|---|---|---|---|
| **Data collection and processing** | | | |
| Magnification | 120,000x | 120,000x | 120,000x |
| Voltage (kV) | 200 | 200 | 200 |
| Electron exposure (e–/Å$^2$) | 40 | 40 | 40 |
| Defocus range (μm) | -0.5 – -2.5 | -0.5 – -2.5 | -0.5 – -2.5 |
| Pixel size (Å) | 1.2 | 1.2 | 1.2 |
| Symmetry imposed | C1 | C1 | C1 |
| Initial particle images (no.) | 1,041,016 | 780,396 | 345,721 |
| Final particle images (no.) | 195,099 | 780,396 | 25,956 |
| Map resolution (Å) FSC = 0.143 | 3.9 | 2.5 | 8.8 |
| Map resolution range (Å) | 2.6 – 16.1 | 2.4 – 4.4 | 5.8 – 13.2 |
| **Refinement** | | | |
| Initial model used | PDB 9NWO | This study | N/A |
| Model resolution (Å) FSC = 0.143 | 4.1 | 2.5 | |
| Model resolution range (Å) | 3.8 – 4.5 | 2.4 – 4.1 | |
| Map sharpening *B* factor (Å$^2$) | -115 | -60 | |
| Model composition | | | |
| Non-hydrogen atoms | 19,342 | 2,138 | |
| Protein residues | 2434 | 266 | |
| Ligands | 4 | 1 | |
| *B* factors (Å$^2$) | | | |
| Protein | 16.38/162.50/73.64 | 9.84/142.88/81.51 | |
| Ligand | 58.85/123.17/95.95 | 93.14/101.48/98.96 | |
| RMSD | | | |
| Bond lengths (Å) | 0.003 | 0.004 | |
| Bond angles (°) | 0.650 | 0.800 | |
| Validation | | | |
| MolProbity score | 1.55 | 1.56 | |
| Clashscore | 3 | 2.38 | |
| Poor rotamers (%) | 0.05 | 0 | |
| Ramachandran plot | | | |
| Favored (%) | 93.21 | 90.46 | |
| Allowed (%) | 6.37 | 9.54 | |
| Disallowed (%) | 0.41 | 0.00 | |
